# Multi-omics studies reveal how ambient temperature changes govern cellular responses of Chlamydomonas

**DOI:** 10.1101/2025.08.21.671244

**Authors:** Prateek Shetty, Trang Vuong, Chulin Li, Volker Wagner, Dinara Myrzakhmetova, Chia-Chi Peng, Wenshuang Li, Joel Ching, Ariane Zander, Sophie Weiser, Miriam A. Rosenbaum, Rosalind J Allen, Markus Lakemeyer, Maria Mittag

## Abstract

Photosynthetic protists, known as microalgae, face increasing temperatures due to climate change. The green biflagellate alga *Chlamydomonas reinhardtii* serves as a model for thermoregulation. While responses to thermal stress are well characterized, much less is known about the impact of ambient temperature shifts. Understanding microalgal responses to environmental temperature changes is critical, as these primary producers drive ecosystem productivity and food web dynamics. Here*, C. reinhardtii* grew mixotrophically at ambient temperatures from 18 °C to 33 °C. Transcriptomic profiling revealed extensive reorganization, with over 5,000 transcripts significantly affected, including those involved in algal-bacterial interactions, photoreception, lipid metabolism, photosynthesis, cilia formation, and the secretome. CO_2_ transfer rates and acetate levels measured at 18 °C and 28 °C suggest decreased photoautotrophic algal growth at 28 °C at first. Antagonistic bacterial activity was sustained longer at lower temperatures. Proteomic analyses of isolated cilia and secreted proteins corroborate major abundance changes within these sub-proteomes, particularly in ciliary intraflagellar transport complexes and mating-related proteins in the secretome. Together, these molecular alterations resulted in pronounced changes in growth, the lengths of cells and cilia swimming behavior, mating ability and bacterial antagonism. These data reveal major cellular responses caused by ambient, even short-term temperature shifts.

## INTRODUCTION

Microalgae contribute significantly to global primary production and are ecologically highly relevant. An often-used model is the unicellular, biflagellate green microalga *Chlamydomonas reinhardtii*. Members of the genus *Chlamydomonas*, including *C. reinhardtii*, have been repeatedly found in flooded rice fields (Barrett and Koch, 1982; Ghasemi et al., 2008; Lin et al., 2013; Carrasco et al., 2024), where acetate is an abundant organic carbon source (Hori et al., 2007).

*C. reinhardtii*, hereafter called Chlamydomonas, is characterized by a single cup-shaped chloroplast, including a pyrenoid along with RUBISCO for carbon dioxide fixation, and a primitive visual system, the eyespot, situated at the edge of the chloroplast near the cell equator (Harris, 1989; Schmidt et al., 2006). Chlamydomonas is a motile alga as it bears two anterior flagella, also called cilia. Each flagellum is anchored by the basal body to the cell. The flagella are structurally and functionally similar to the 9+2-type axonemes of most motile mammalian cilia (Pazour et al., 2005). The flagella contain an intraflagellar transport system (IFT) that is situated between the axoneme and the ciliary membrane and connected with flagellar length control (Marshall and Rosenbaum, 2001). The ciliary axoneme consists of substructures such as dynein arms and radial spokes that control bending and are essential for motility, mating, and environmental sensing (Marshall, 2024).

The nuclear and organelle genomes of Chlamydomonas are fully sequenced and annotated. In addition, its defined sexual breeding cycle involving plus and minus gametes, along with extensive mutant collections of haploid vegetative cells, allows for genetic crosses and offers precise genetic manipulation using existing toolkits (Merchant et al., 2007; Crozet et al., 2018; Li et al., 2019; Craig et al., 2023). These features have firmly established Chlamydomonas as a model organism for studying a variety of biological processes such as photosynthesis, light perception, cilia biogenesis or synthetic biology (Scaife and Smith 2016; Dupuis and Merchant, 2023).

In recent years, algal-bacterial interactions have come into focus. Algae and bacteria coevolved for hundreds of millions of years, fostering highly relevant ecological associations (Burgunter-Delamare et al., 2024). Chlamydomonas also emerged as a model to study bacterial interactions. Examples include the vitamin B_12_-based mutualism (Helliwell et al., 2015; Bunbury et al., 2022) or the antagonism of the bacterium *Pseudomonas protegens* Pf-5 (Aiyar et al., 2017). *P. protegens* secretes toxins including the cyclic lipopeptide orfamide A that deflagellates the cell within a minute by increasing the algal cytosolic Ca^2+^ levels. At least four different algal transient receptor potential channels (TRPs) are involved in mediating the orfamide A signals (Hou et al., 2023).

Temperature is a critical factor in the life of many species. The effects of changes in ambient temperature on microalgal life are, however, poorly understood. Climate warming shifts the species distribution of animals, plants and phytoplankton (Mäkinen et al., 2025) and thus can influence ecosystems. Ambient temperature changes may influence the growth and metabolism of algae, while extreme temperatures may induce stress responses in the short term and lead to cell death in the long term. With climate change driving temperature fluctuations, understanding cellular responses to ambient temperature changes in model algae like Chlamydomonas is essential. Chlamydomonas is conventionally cultivated in the range of 20 °C to 25 °C in the laboratory (Harris, 1989). The alga begins to experience heat stress above 37 °C, marked by the activation of *heat shock protein* (*HSP*) genes, and experiences strong heat stress when the cultivation temperature approaches 43.5 °C (Schroda et al., 2015). A lipidome analysis revealed that Chlamydomonas cells exposed to heat stress of 42 °C exhibited changes in lipid metabolism and increased lipid saturation as an early heat response (Légeret et al., 2016). A recent system-wide analysis made use of transcriptomics and proteomics to study the response of Chlamydomonas to moderate (35 °C) temperature stress that transiently arrests the cell cycle and acute temperature stress (40 °C) disrupting cell division and growth (Zhang et al., 2022). The results showed that some of the responses were shared between the two treatments, although the effects of the 40 °C stress were typically more extensive. It was also found that cultures treated at 35 °C increase their consumption of acetate (Zhang et al., 2024). Besides high temperatures, temperatures below 13 °C can induce a cold stress response in Chlamydomonas. Cold stress causes the arrest of cell division, increases the cell size and induces palmelloid formation (Ermilova, 2020). Low temperature adaptation in microalgae also involves production of betaine lipids that are nonphosphorous polar glycerolipids (Murakami et al., 2018) and are specifically found in some microalgae including Chlamydomonas (Riekhof et al., 2005).

Here, we aimed to investigate primarily changes in ambient temperature over several diurnal cycles. In our study, Chlamydomonas was maintained under non-stress temperatures including 18 °C, 23 °C, 28 °C, and 33 °C for four days, allowing system-wide profiling of its adaptive responses during active growth. The algal cells were grown in a medium with acetate, as found in one of its natural habitats.

Our comprehensive analysis integrates results from multiple assays across the four above-mentioned ambient temperatures. Transcriptomic analysis reveals how temperature modulates the expression of transient receptor potential (TRP) channels that are important for some algal-bacterial interactions, as well as components of RNA and lipid metabolism, photosynthesis, major photoreceptors, and numerous ciliary and secretome-related transcripts. Based on growth data, we found that increasing temperatures promote algal growth with an optimum around 28 °C. Our transcriptomic results are extended by immunoblots of selected photoreceptors, analyses of the ciliary and secretory proteomes as well as by behavioral assays. Elevated temperatures upregulate many secreted proteins, including those related to mating. A significant increase in gamete mating efficiency was found. Furthermore, proteins of the IFT complexes associated with flagellar length control, as well as other key ciliary structural proteins, become more abundant at 28 °C, whereas acetylated α-tubulins and α-tubulins decrease. Importantly, morphological examinations reveal that the lengths of cells and cilia decrease in cells grown at higher ambient temperatures. Swimming behavior is also affected by ambient temperatures. At 28 °C, algal cells swim at a reduced speed compared to 18 °C. Moreover, the swimming path is characterized by many changes in direction at 28 °C. The frequent change in swimming direction at 28 °C is independent from acclimation, as it is triggered within just 15 min of a temperature shift. Together, these results demonstrate that thermal effects, including acclimation at different ambient temperatures, impact not only the transcriptome and proteome in Chlamydomonas, but also profoundly influence cell morphology, interspecies interactions, motility, and the sexual cycle.

## RESULTS

### Ambient temperatures influence the growth and cell length of Chlamydomonas and shape its transcriptome

To address how Chlamydomonas may modulate its physiology and gene expression under non-stressful, ecologically relevant temperature settings, we systematically profiled growth, cell length, and global transcript abundances across a spectrum of ambient temperatures (18 °C, 23 °C, 28 °C, and 33 °C). Cells were grown in a light-dark cycle of 12 h light and 12 h dark (LD 12:12) in acetate-containing TAP medium and investigated during the day (see Methods). As mentioned before, acetate is also present in the natural environment of Chlamydomonas in rice fields and enables mixotrophic algal growth. Chlamydomonas cells are synchronized under mixotrophic conditions in a LD cycle using acetate as carbon source (Catalan et al., 2025).

Our results show that cultivation at 18 °C results in a significantly lower growth rate and reduced final cell density compared to the usual growth temperature of 23 °C, while elevated cultivation temperatures of 28 °C and 33 °C accelerated early growth and achieved greater final biomass (Figure 1A, Supplementary Table 1). The growth limitation at 18 °C becomes evident early in cultivation, with significant differences (compared to growth at the reference temperature of 23 °C) being observed consistently from day 3 onwards, ultimately resulting in a significantly reduced final cell density of 1.20 × 10^7^ cells mL^-1^. In contrast, cultures grown at 28 °C achieved a final cell density of 1.84 × 10^7^ cells mL^-1^, representing an approximately 20% increase over the 23 °C reference (1.54 × 10^7^ cells mL^-1^). Although the 33 °C cultivation yielded a slightly higher final cell density (1.88 × 10^7^ cells mL^-1^ at day 15) than the 28 °C cultivation, cell densities from days 3 to 14 were in most cases higher at 28 °C, indicating that the optimal temperature range for growth is around 28 °C (Figure 1A, Supplementary Table 1).

**Figure 1:**
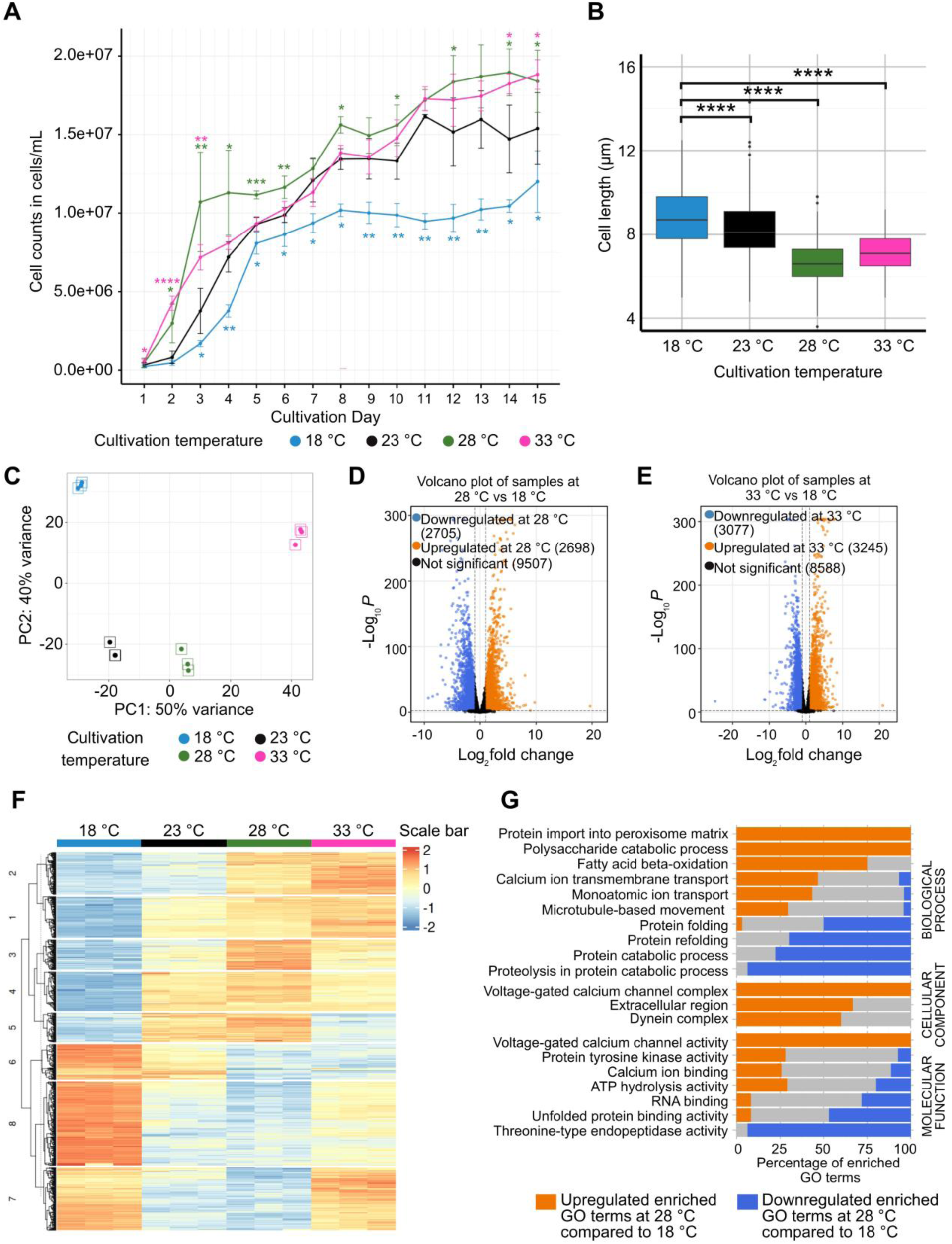
Influence of different ambient cultivation temperatures on the growth and transcript abundances of Chlamydomonas. **A)** Growth curve of Chlamydomonas cultured at 18 °C (blue), 23 °C (black), 28 °C (green), and 33 °C (magenta) over a 15-day period. Data points represent mean cell densities (cells mL^-1^) from two independent datasets, each comprising three biological replicates (Supplementary Table 1). Exceptions are days 8 and 11 comprising three biological replicates solely. For better visibility, individual data points of these algal monocultures for days 8 and 11 have been inserted in Figure 2C (18 °C), Figure 2D (28 °C) as well as Supplementary Figures 5A (23 °C) and 5B (33 °C) as open circles. Error bars indicate standard deviation. Asterisks indicate the statistically significant differences for 18 °C (blue), 28 °C (green), and 33 °C (magenta) compared to 23 °C (black) as calculated by a two-sided *t*-test assuming unequal variance (Welch’s *t*-test) (*: *P* < 0.05, **: *P* < 0.01, ***: *P* < 0.001, ****: *P* < 0.0001). **B)** Boxplot of vertical cell length measurements (µm) on day 4 of cultivation across the four temperature treatments. The cell lengths of day 1 and day 7 are shown in Supplementary Figure 1A, B. Each box represents measurements from three biological replicates (100 cells per replicate). Asterisks represent significant differences as calculated by a two-sided *t*-test assuming unequal variance (Welch’s *t*-test; ****: *P* < 0.0001). **C)** Principal component analysis (PCA) of transcriptomic profiles from Chlamydomonas after 4 days of cultivation at each temperature. Each point corresponds to a biological replicate, colored by temperature (18 °C, blue; 23 °C, black; 28 °C, green, and 33 °C, magenta). PC1 and PC2 account for 50% and 40% of the total variance, respectively, illustrating clear separation by temperature, especially at the extreme temperature conditions. **D, E)** Volcano plots depicting differentially abundant transcripts between 18 °C and 28 °C (D), and 18 °C and 33 °C (E), respectively. Transcripts with significant upregulation (orange) and downregulation (blue) are defined by adjusted *P*-value < 0.01 and |Log_2_ FC| ≥ 1. Non-significantly changed transcripts are shown in black. Total transcript counts for each category are indicated in parentheses. Vertical dashed lines mark log_2_ fold change thresholds of ±1; the horizontal dashed line marks the significance threshold. **F)** 1000 differential transcript abundances from cells grown either at 18 °C or 28 °C were determined. Transcripts displaying the most dominant abundances, being either upregulated (500 transcripts) or downregulated (500 transcripts) were clustered across the four temperatures in a heatmap. Each row represents a gene, and each column a temperature. **G)** Exemplary results from a Gene Ontology (GO) enrichment analysis of differentially expressed genes, comparing 18 °C and 28 °C. The full list is shown in Supplementary Table 4. Bar plots display the percentage of enriched GO terms in Biological Process, Cellular Component, and Molecular Function categories. Orange bars indicate GO terms enriched at 28 °C relative to 18 °C, while blue bars represent downregulated terms.

We also found that the vertical cell length of the algal cells, measured in the middle of the day at LD6 (Figure 1B), was significantly reduced upon elevation of the cultivation temperature. On day 4, cells grown at 18 °C averaged 8.80 µm in length. Cells cultivated at 23 °C exhibited a modest yet significant decrease in mean cell length (to 8.25 µm). Further increasing the temperature to 28 °C and 33 °C intensified this effect, causing the cells to shrink to 6.65 µm at 28 °C and 7.15 µm at 33 °C, respectively (Figure 1B, Supplementary Table 2A). The observation that temperature-dependent effects are slightly more pronounced at 28 °C versus 33 °C suggests that the minimal cell length has already been reached at 28 °C. Similar data were obtained when cell lengths were determined as early as day 1 of cultivation (Supplementary Figure 1A) and later on day 7 for cultures grown at the higher temperatures compared to 18 °C (Supplementary Figure 1B). Data measurements were performed in the middle of the day (LD6) on day 1 pointing to a relatively direct heat-effect. We also determined the cell length distribution pattern at different temperatures at LD6 to assess the synchrony of cells on day 1 to day 4. The pattern was found to be unimodal in all cases (Supplementary Figure 2).

To approximate in addition cell areas, we manually traced individual cell boundaries using the Polygon Selection tool for days 1 and 4. These estimated cell area measurements support our vertical cell length data (Supplementary Figure 1C, D; Supplementary Table 2B). Specifically, we observed a similar trend where higher temperatures resulted in significantly smaller cell areas.

In the next step, we performed comprehensive RNA-seq profiling of cells grown for four days at temperatures of 18 °C, 23 °C, 28 °C, and 33 °C, with three biological replicates at each temperature. Our results revealed marked and temperature-dependent shifts in the global Chlamydomonas transcriptome (Supplementary Table 3). All twelve libraries exceeded 58 million raw reads, with ≥83% of trimmed reads aligning to the Chlamydomonas reference genome v6.1. The average mapping rate ranged from ∼83% to ∼87%. Principal-component analysis (PCA) separated all four temperature conditions along PC1 (50% variance) and PC2 (40% variance), indicating broad transcriptional reprogramming with minimal between-replicate variation (Figure 1C). Together, PC1 and PC2 captured 90% of the total variance. Differential expression analysis highlighted that thousands of transcript abundances were either increased or decreased at 28 °C and 33 °C, relative to 18 °C (Figure 1D, E), eliciting large-scale transcriptional remodeling. A total of 5,403 genes at 28 °C and 6,322 genes at 33 °C, respectively, were found to be differentially abundant at a |log₂ fold change (FC)| ≥ 1 and adjusted *P*-value < 0.01 relative to 18 °C.

Growth measurements indicated that Chlamydomonas grows better as temperature increases from 18 °C, with the optimal growth being at 28 °C and no further gains being observed above this temperature (Figure 1A). Thus, we focused our subsequent analysis on the results from the 18 °C versus 28 °C comparison on day 4. To visualize the most dominant transcriptional patterns in a heatmap, the top 1,000 differentially expressed transcripts from this comparison were selected, comprising (i) the 500 transcripts with the largest positive log_2_ FC and (ii) the 500 transcripts with the largest negative log_2_ FC (Figure 1F). These genes were filtered from the full list of statistically significant transcripts (adjusted *P*-value < 0.01, |log_2_ FC| ≥ 1) to highlight the dominant transcriptional signatures across all temperature conditions. A heatmap of all differentially abundant transcripts confirms these dominant temperature-dependent transcript patterns (Supplementary Figure 3).

Gene ontology (GO) enrichment of the transcripts that were differentially abundant between the 18 °C and 28 °C samples revealed several functionally important categories (Figure 1G, Supplementary Table 4). Extracellular matrix (ECM) remodeling emerges as an important theme, with upregulated GO terms including “extracellular region (GO:0005576)” and “polysaccharide catabolic process (GO:0000272)”. The Chlamydomonas ECM is composed of pherophorins and other hydroxyproline-rich glycoproteins (HRGPs) and is closely connected to the secretome (Luxmi et al. 2018). Based on these results, it seems that the ECM may undergo modification under different ambient cultivation temperatures. Additionally, flagellar-related GO terms show upregulation at 28 °C, including “dynein complex (GO:0030286)”, “ATP hydrolysis activity (GO:0016887)”, and “protein tyrosine kinase activity (GO:0004713)”. The increased expression of dynein motor proteins and kinase activities suggests enhanced flagellar motor activity. Besides, enrichment of “ATP hydrolysis activity” may be related to the fact that ciliary motility is driven by dynein motors that hydrolyze ATP (Mitchell et al., 2005). The upregulation of “protein import into peroxisome matrix (GO:0016558)” could indicate altered protein trafficking pathways that may affect flagellar protein assembly. Moreover, protein folding processes including “protein folding (GO:0006457)”, “protein refolding (GO:0042026)”, and “unfolded protein binding (GO:0051082)” are downregulated at 28 °C.

### Transcripts encoding photoreceptors and TRP channels show temperature-dependent regulation along with an altered bacterial antagonism against Chlamydomonas

TRP channels such as the human TRPA1 (Moparthi et al., 2022) or TRP1 of Chlamydomonas (McGoldrick et al., 2019) are known for sensing temperature. Several algal TRPs participate in controlling algal-bacterial interactions (Hou et al., 2023). Transcriptomic profiling identified robust temperature-dependent regulation of transcripts encoding TRP channels (Figure 2A, Supplementary Figure 4). 18 of the 29 known *TRP*s (Hou et al., 2023) display significant differential abundance at 28 °C or 33 °C compared to 18 °C with a |log₂ FC| ≥ 1 in their abundances (Supplementary Figure 4, Supplementary Table 5). *TRPs 34* and *5* transcripts are the most strongly downregulated at 33 °C compared to 18 °C (log_2_ FC close to –11 and –8, respectively). Interestingly, three known *TRP* transcripts implicated in the orfamide A-driven signaling process, *TRP*s 5, 11, and 22 (Hou et al., 2023) are altered in their abundance. Besides the *TRP5* channel transcript, *TRP11* also exhibits significant repression at elevated temperatures, while *TRP22* is slightly enhanced. These changes in the expression of *TRP*s that are involved in orfamide A perception suggest that higher temperatures may alter the vulnerability of Chlamydomonas to orfamide A-mediated bacterial attacks. To test if there was any temperature-dependent change of bacterial antagonism, we cocultured Chlamydomonas with *P. protegens* at a ratio of 1:250, previously shown to be efficient for the bacterial attack (Carrasco et al., 2024) in TAP medium at 18 °C, 23 °C, 28 °C, and 33 °C, respectively. Inspection of the green color of the cultures was used to estimate bacterial antagonism and recovery of the algal cells (Figure 2B). We observed that the algal cells in algal-bacterial cocultures cultivated at higher ambient temperatures (either 28 °C or 33 °C) apparently recovered by day 8 of growth in coculture, while they had not recovered by this time at lower ambient temperatures (Figure 2B). To track algal cell density more quantitatively, we counted algal cell numbers in monocultures and in cocultures together with *P. protegens*. At 18 °C, Chlamydomonas in coculture with *P. protegens* exhibits reduced growth, with no or little recovery until 11 days (Figure 2C). Similar results are obtained for a coculture grown at 23 °C (Supplementary Figure 5A). In contrast, at 28 °C and 33 °C, algal cells exhibit a much faster recovery in cocultures, resuming growth by day 7 and achieving cell densities closer to those of axenic controls by day 15 (Figure 2D, Supplementary Figure 5B). We also analyzed the growth of *P. protegens* at the lower (18 °C) compared to higher temperatures (28 °C and 33 °C) until recovery day 8. Like the algae, the bacteria grow faster at higher temperatures (Figure 2E, Supplementary Table 6). Notably, they reach their optimum growth on day 1 to 2 at the higher temperatures and then decline in density until day 8 to a rather low level. In comparison, bacteria grown at 18 °C are at a significantly higher optical density on day 8, which agrees with the longer duration of their antagonism.

**Figure 2:**
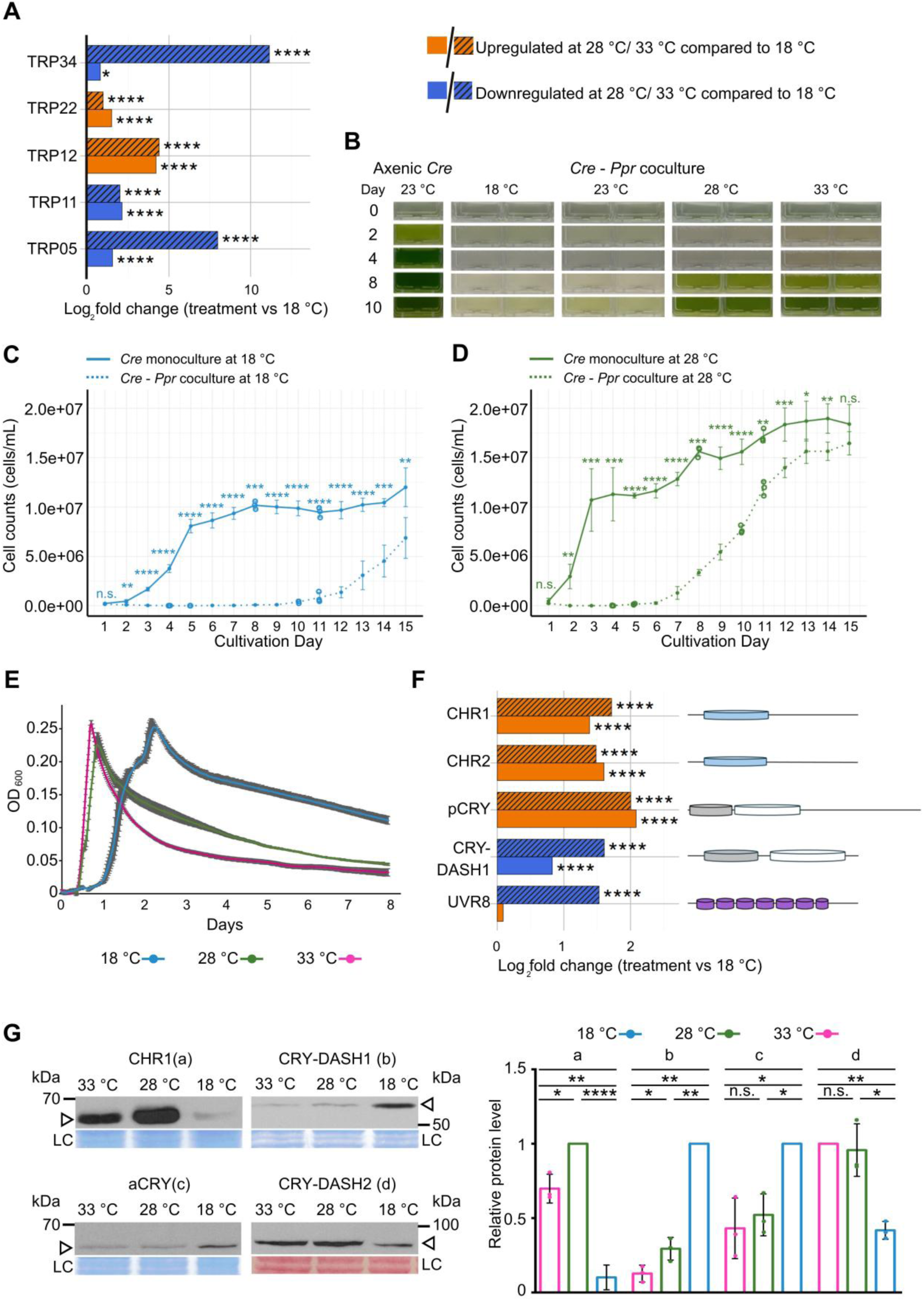
Temperature-dependent regulation of transcripts encoding selected TRP channels and photoreceptors along with mitigation of bacterial antagonism. **A)** Log_2_ fold changes in the expression of selected *TRP* channel transcripts. A list of all significantly changed *TRP* channel transcripts with a |log₂ FC| ≥ 1 in Chlamydomonas in response to cultivation at varying temperatures (18 °C, 28 °C, and 33 °C) is presented in Supplementary Figure 4 and Supplementary Table 5. TRPs 5, 11, and 22 are known to be involved in the orfamide A-triggered signaling response (Hou et al., 2023). **B)** Visual evidence indicating the attenuation of bacterial antagonism during cocultivation of Chlamydomonas (abbreviated as *Cre*) with *P. protegens* (*Ppr)* at higher temperatures. Green chlorophyll coloration at 28 °C and 33 °C denotes improved algal viability compared to 18 °C, indicating that increased temperature diminishes bacterial antagonistic effects. **C, D)** Growth curves of axenic *Cre* (monocultures) and *Cre* - *Ppr* cocultures at 18 °C and 28 °C, respectively. Cell densities (cells mL^-1^) were measured over 15 days, highlighting temperature-associated changes in antagonism. Data points represent mean ± standard deviation cell densities from two independent datasets, each comprising three biological replicates, unless otherwise indicated. Data from *Cre* monocultures have been taken from Figure 1A (Supplementary Table 1). In the *Cre* - *Ppr* cocultures, only three biological replicates are available on certain days. Specifically, biological replicates are missing for days 5 and 11 of dataset 1 and for days 4 and 10 in dataset 2. Individual data from the three present biological replicates from days 4, 5, 10 and 11 are indicated by open circles. Statistical significance is indicated by asterisks. **E)** Growth curves of *Ppr* monocultures grown at 18 °C, 28 °C, and 33 °C. Cell densities (OD_600_) were measured over 8 days (Supplementary Table 6), using a plate reader. Data points represent mean ± standard deviation cell densities from three biological replicates. Each biological replicate includes eight technical replicates. **F)** Log_2_ fold changes (|log₂ FC| ≥ 1) in the expression of photoreceptor transcripts (Supplementary Table 5) under varying temperatures. Known domains of the photoreceptors analyzed by Phytozome 14 (https://phytozome-next.jgi.doe.gov/) are highlighted (Greiner et al., 2017). Blue colored circles indicate bacterial/fungal rhodopsin-like protein domains, grey circles DNA-photolyase domains with antenna chromophores MTHF (5,10- methenyltetrahydrofolate) or 8-HDF (8-hydroxydeazaflavin), white circles FAD binding domains of DNA-photolyases and violet circles regulator of chromosome condensation (RCC) repeats. For simplicity, the size of the proteins has been standardized to a common bar length; the relative position of the domains is given. The bar size for plant cryptochrome (pCRY) with more than 1,000 amino acids has been enlarged. CHR, channelrhodopsin; CRY-DASH, *Drosophila Arabidopsis Synechocystis* human cryptochrome; UVR8, UV receptor8. **G)** Left panel: Immunoblot analysis showing CHR1 (a), CRY-DASH1 (b), aCRY (c) and CRY-DASH2 (d) protein abundances in cells grown at 18 °C, 28 °C, and 33 °C. 75 µg of total protein per lane were separated by SDS-PAGE and immunoblotted with the respective antibodies as described in the Methods section. n = 3 biological replicates. In case of CHR1 (cells grown at 18 °C or 28 °C) six biological replicates were performed. Arrows indicate the positions of the target protein. Selected bands from a polyvinylidene difluoride membrane or a nitrocellulose membrane (for anti-CRY-DASH2 blots) stained with Coomassie Brilliant Blue R250 or Ponceau (for anti-CRY-DASH2 blots) were used as a loading control (LC). Right panel: Quantification of immunoblots. The highest level of each protein at any of the three temperatures was always set to 1. **C, D, G**) Asterisks represent significant differences as calculated by a two-sided *t*-test assuming unequal variance (Welch’s *t*-test). *P* < 0.05 (*), *P* < 0.01 (**), *P* < 0.001 (***), and *P* < 0.0001 (****). **A, F**) Adjusted *P*-values < 0.01 (*) and < 0.00001 (****), as calculated by DESeq2, are shown.

Besides TRPs, photoreceptors are known to sense temperatures, at least in land plants (Paik and Huq, 2019). Our analysis revealed temperature-dependent reprogramming of five of the 20 predicted/known photoreceptor transcripts (Figure 2F; Greiner et al., 2017; Sharma et al., 2025). Transcripts encoding channelrhodopsins (CHR1, CHR2), which are located in a specialized part of the plasma membrane covering the eyespot area display pronounced upregulation at higher temperatures (Figure 2F). Conversely, one of the four known cryptochrome transcripts, CRY-DASH1, shows a strong decrease in its abundance as the temperature increases. We examined whether the strong up- and down-regulation of the CHR1 and CRY-DASH1 transcripts, respectively, is reflected at the protein level, using immunoblot analyses. For CRY-DASH1, the protein levels mirrored exactly the tendency found in the transcript profile at 18°C, 28 °C, and 33 °C with a peak at 18 °C (Figure 2G). In case of CHR1, the protein level was increased at both higher temperatures but reached a maximum at 28 °C, while the transcript was most abundant at 33 °C. We also checked another photoreceptor, the animal-like cryptochrome (aCRY) whose transcript level does not exhibit variations with a |log₂ FC| ≥ 1. It was shown recently that aCRY can be controlled at the post-transcriptional/translational level (Vuong et al., 2025). aCRY can indeed sense ambient temperature; its protein amount is upregulated at 18 °C compared to 28 °C or 33 °C (Figure 2G). Finally, we examined CRY-DASH2 (Rredhi et al., 2021) whose function is still open. In contrast to CRY-DASH1, CRY-DASH2 is strongest upregulated at 33 °C compared to 18 °C (Figure 2G), opening interesting perspectives for its function (see discussion).

As posttranscriptional/translational regulation seems involved in some cases, we screened RNA-binding proteins (RBPs), which tune gene expression post-transcriptionally or translationally. Nine transcripts encoding RBPs are responsive to ambient temperatures (Supplementary Figure 6). Some RBP transcripts, encoding Pumilio or Musashi proteins, are mostly significantly upregulated at higher temperatures, while the C3 and C1 subunit transcripts of the RNA-binding protein CHLAMY1 (*CRB1* and *CRB3*; Zhao et al., 2004) as well as NAB1 and DUS16 are downregulated at higher temperatures. Some of these data are reflected at the protein level, as discussed later.

### Ambient temperature drives changes in transcripts encoding proteins of the ciliary IFT, the secretory and several metabolic pathways thereby controlling CO_2_ consumption

To further elucidate the molecular adaptations underlying temperature-dependent phenotypic changes in Chlamydomonas, we investigated the expression of additional key gene families highlighted by the gene ontology enrichment, focusing on flagellar assembly, ECM remodeling and enzymes involved in metabolic pathways (Supplementary Tables 5 and 7).

Flagellar trafficking systems, specifically the IFT complexes, are essential for the assembly, maintenance, and function of eukaryotic flagella (Pedersen et al., 2006; Katoh et al., 2016; Ma et al., 2023). The IFT is composed of IFT-A and IFT-B complexes, which coordinate the bidirectional transport of structural and signaling components along the axoneme. Proper IFT activity is fundamental for Chlamydomonas motility, environmental sensing, and cell division. In our transcriptome analysis (Figure 3A), we observed robust and coordinated upregulation of one *IFT-A* and many *IFT-B* subunit transcripts in cells grown at both 28 °C and 33 °C compared to 18 °C. These findings, if reflected at the protein level, imply that the upregulation of IFT machinery at higher temperatures reflects a state of active flagellar remodeling.

**Figure 3:**
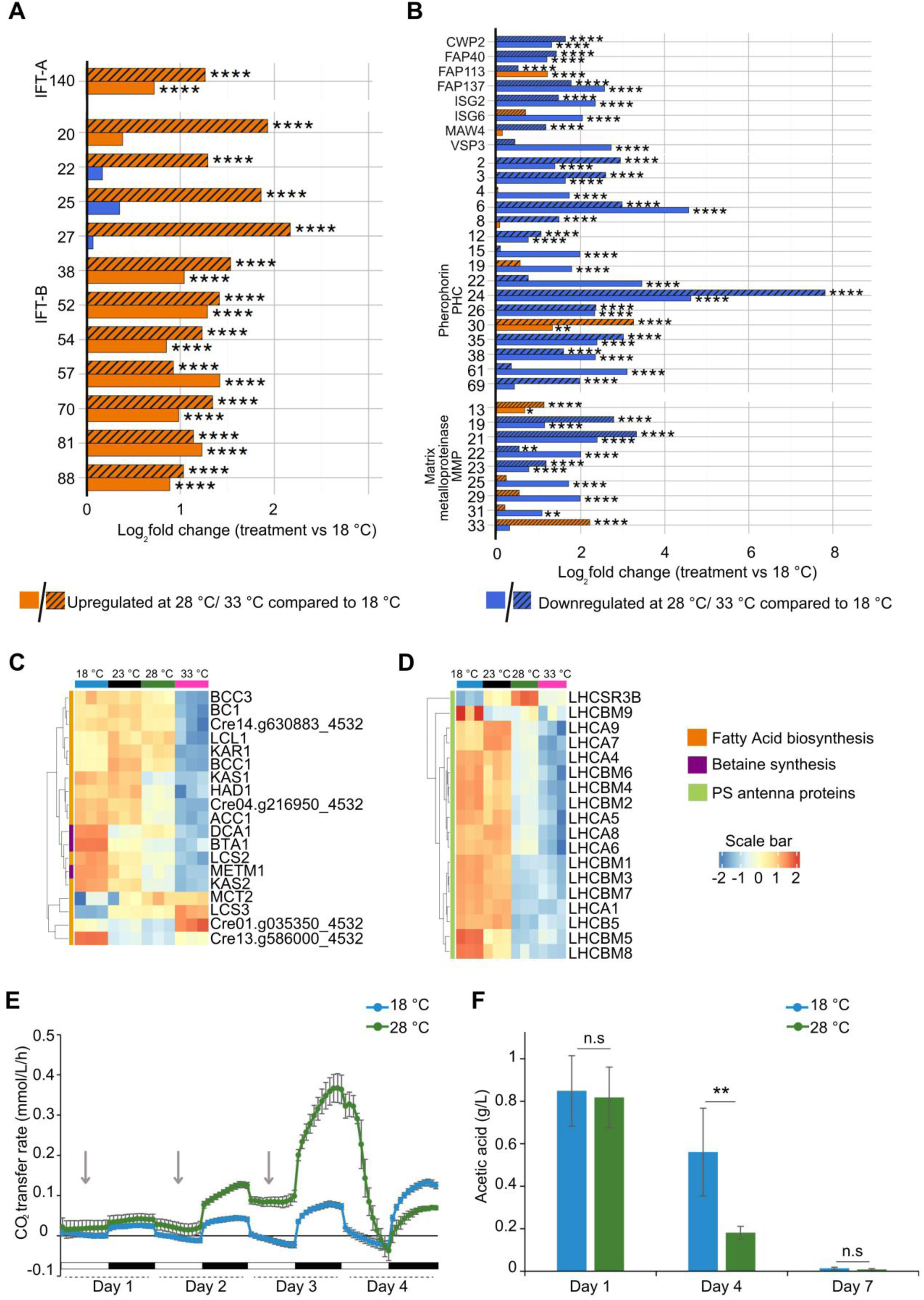
Temperature-dependent regulation of transcripts encoding IFT and extracellular proteins as well as metabolic proteins along with photoautotrophic growth and acetate availability. **A)** Expression of transcripts encoding IFT proteins. Several transcripts encoding IFT-A and IFT-B complex proteins with a |log₂ FC| ≥ 1 show significantly higher abundance at temperatures of 28 °C and/or 33 °C. **B)** Expression of transcripts with a |log₂ FC| ≥ 1 encoding predicted extracellular proteins as identified by DeepLoc (see Methods) in cells grown at different temperatures. The first eight transcripts encode hydroxyproline-rich glycoproteins. CWP2, cell wall protein2, also known as FAP3; FAP, flagellar associated protein; ISG2/6, inversion-specific glycoprotein, found in cilia; MAW4, membrane-associated wall protein; VSP3, serine-proline repeat hydroxyproline-rich cell wall protein found in cilia. **A, B)** Adjusted *P*-values < 0.01 (*), < 0.001 (**), and < 0.00001 (****), as calculated by DESeq2, are shown. **C)** Heatmap depicting expression of transcripts with a |log₂ FC| ≥ 1 involved in the fatty acid biosynthesis pathway. BCC3 (acetyl-CoA biotin carboxyl carrier), BC1 (biotin carboxylase/ACCase complex), Cre14.g630883_4532 (K03921 - acyl-DESA1), LCL1 (long-chain-fatty-acid-CoA ligase), KAR1 (3-oxoacy-ACP reductase), BCC1 (acetyl-CoA biotin carboxyl carrier), KAS1 (3-ketoacyl-ACP-synthase), HAD1 (3-hydroxyacyl-ACP dehydratase), Cre04.g216950_4532 (putative 3-ketoacyl-ACP-synthase), ACC1 (ACCase complex), DCA1 (S-adenosyl-L-methionine decarboxylase), BTA1 (betaine lipid synthase), LCS2 (long-chain acyl-CoA synthetase), METM1 (S-adenosylmethionine synthase), KAS2 (3-ketoacyl-ACP-synthase), MCT2 (malonyl-CoA:acyl-carrier-protein transacylase), LCS3 (long-chain acyl-CoA synthetase), Cre01.g035350_4532 (K07512 - mitochondrial trans-2-enoyl-CoA reductase), Cre13.g586000_4532 (K18660 malonyl-CoA) **D)** Heatmap depicting expression of transcripts encoding PSII antenna proteins with a |log₂ FC| ≥ 1. LHC (light-harvesting complex). (**A-D**) Genes/proteins are listed in Supplementary Table 5 along with their Cre accession numbers. **E)** CO_2_ consumption and production in cells grown in a LD cycle in TAP medium at 18 °C and 28 °C, respectively, over four days. Data were measured every 60 min. Arrows highlight the first three-day phases of the LD cycle. Three biological replicates were performed (see Methods). For better visibility, individual data points of all replicates from the four days measured every 60 min have been listed in Supplementary Table 8. **F)** Acetate content in cultures grown in an LD cycle in TAP medium at 18 °C and 28 °C, respectively, during day 1, day 4 and day 7 of growth. The experiments were performed using two independent datasets, each along with three biological and technical replicates (see Methods). Asterisks represent significant differences as calculated by a two-sided *t*-test assuming unequal variance (Welch’s *t*-test). *P* < 0.01 (**), n.s., not significant.

Given the enrichment of ECM-related terms, we next examined changes in transcripts encoding ECM proteins. The ECM in Chlamydomonad algae such as *Volvox carteri* is primarily composed of hydroxyproline-rich glycoproteins, particularly pherophorins and related structural proteins. Phylogenetic analyses comparing *V. carteri* and Chlamydomonas have revealed that both species possess numerous pherophorin-related proteins. In *V. carteri*, 74 of 99 of these proteins are species-specific, while in Chlamydomonas, 10 of 30 are unique to the species (von der Heyde and Hallmann, 2022). These ECM components play a pivotal role in maintaining cell wall integrity, protection, signaling, and morphogenesis. They not only provide mechanical stability but also participate in cell-cell communication and environmental adaptation (Hallmann, 2006). Matrix metalloproteases (MMPs) are key regulators of ECM remodeling, cleaving glycoproteins to facilitate matrix turnover and structural adaptation. Both are tightly connected to the algal secretome (Luxmi et al., 2018). Our data show consistent and dramatic downregulation of multiple transcripts encoding ECM proteins and MMPs at higher temperatures (Figure 3B).

We also addressed metabolic pathways. While glycolysis and the TCA cycle do not show a coordinated transcriptional response to ambient temperature changes (Supplementary Table 7), lipid biosynthesis demonstrates a coordinated response as visualized in a heatmap (Figure 3C; Supplementary Table 5). Most transcripts encoding enzymes of fatty acid biosynthesis are downregulated at 33 °C. Such a downregulation also includes betaine lipid synthesis, which is known to be involved in abiotic stresses, including cold stress (Salomon et al., 2025).

We further examined temperature-dependent regulation of transcripts encoding proteins of photosynthesis. Transcripts encoding the antenna proteins of the photosystems are mostly upregulated at lower temperatures (Figure 3D) as well as proteins of photosynthesis and enzymes involved in chlorophyll biosynthesis (Supplementary Figure 7, Supplementary Table 5), while transcripts encoding enzymes of the Calvin cycle do not exhibit a coordinated response (Supplementary Table 7). However, CO_2_ availability in solution is usually higher at lower temperatures and is likely to affect the key CO_2_ fixing activity of the enzyme RUBISCO. We examined the effect of CO_2_ availability in pure water at 18 °C compared to 28 °C and 33 °C (Supplementary Figure 8). The CO_2_ concentration drops by about one third from 18 °C to 28 °C and 33 °C, respectively. Taken together, these findings raise the question of whether photoautotrophic growth is favored at 18 °C relative to 28 °C, given that acetate supports heterotrophic growth in algae. As the cells were cultured in TAP medium, they grow mixotrophically and may preferentially metabolize either CO₂ or acetate. To get more insight into photoautotrophic growth at both temperatures, we compared CO_2_ transfer rates in cells grown in a LD cycle in TAP medium at 18 °C versus 28 °C from day 1 to 4 with the help of an incubation chamber connected to an automated CO_2_/O_2_ off-gas analysis (see Methods). The CO_2_ transfer rate (CTR) was recorded every 60 min (Figure 3E, Supplementary Table 8). Conventionally, CO_2_ will be consumed during the day by algal photosynthesis and produced during the night via respiration, leading to negative and positive CTRs, respectively. In cells grown at 18 °C, the CTRs follow exactly this pattern. Thus, CO_2_ levels decline during the day and increase during the night during the four days. In contrast, cells grown at 28 °C reveal a clearly different pattern. CO_2_ consumption during the day is rather constant in these cells during the first three days (indicated by grey arrows), even though cell density is higher at 28 °C compared to cells grown at 18 °C (Figure 1A). The higher cell number at 28 °C is well reflected by the higher CO_2_ production at night at 28 °C. We conclude that cells grown at 18 °C rely mainly on photoautotrophic growth during the first three days, while cells grown at 28 °C seem to grow rather heterotrophically using acetate during this time. Only at day 4, when the amount of produced CO_2_ by respiration during the night is at a very high rate, which is likely to be reflected by higher CO_2_ levels in solution, cells grown at 28 °C start to consume CO_2_ at a high rate during the day. The differences in photoautotrophic growth change could be additionally related to the potential absence of acetate available on day 4. We measured the acetate levels in the culture medium of cells grown at 18 °C and 28 °C, respectively, on day 1, 4 and 7 (Figure 3F). On day 1, there is no significant difference between the level of acetate in cultures grown at 18 °C versus 28 °C. These levels are similar to those in fresh TAP medium (about 1 g/L). In contrast, on day 4, acetate levels are significantly reduced, especially in the culture medium of cells growing at 28 °C, but are still in a sufficient range. Acetate is thus not missing in cultures grown at 18 °C and 28 °C, respectively, on day 4 (Figure 3F). In contrast, acetate is used up at day 7 in cultures grown at 18 °C and 28 °C (Figure 3F). The very high CO_2_ levels produced in the night phase of day 3 at 28 °C may contribute to switching these cells from mainly heterotrophic growth to photoautotrophic growth on day 4. Our data emphasize that the ratio of photoautotrophic versus heterotrophic growth of Chlamydomonas cells grown mixotrophically in acetate-containing medium is altered upon changes in ambient environmental temperature.

### Temperature-dependent remodeling of the flagellar proteome results in shorter flagella in cells grown at 28 °C

Transcriptome analysis revealed that many transcripts encoding ciliary IFT proteins are altered in their abundances in cells grown at 18 °C compared to 28 °C (Figure 3A). To examine whether these changes are reflected at the protein level, we performed label-free quantitative proteomic analyses on ciliary proteins of purified NP-40 treated flagella from cells grown at 18 °C or 28 °C (see Methods) using single-pot phase-enhanced sample preparation (Hughes et al., 2019) and liquid chromatography electrospray ionization tandem mass spectrometry (LC-ESI-MS/MS, see Methods). To ensure high-quality proteome data, we applied a 2.5% peptide count filter based on the maximum peptide count per protein for the dataset (see Methods). With a maximum of 304 unique peptides per protein in the ciliary proteome, a protein requires at least eight unique peptides to qualify. By this way, we identified a total of 1,109 proteins with at least eight unique peptides (Supplementary Table 9). We generated a high-confidence ciliary protein list by matching all detected proteins against the Chlamydomonas ciliary protein database (see Methods) and labeled these candidates in dark blue in Supplementary Table 9. Moreover, we performed a manual inspection of all other candidates against existing ciliary databases (see Methods) and current literature. We provide a reference for each of the identified candidates in Supplementary Table 9. These manually verified ciliary candidates are indicated in light blue in Supplementary Table 9, while proteins of the manual inspection with unknown or overly broad function are marked in red. Potential contaminants are denoted by a white background.

Through this pipeline, we identified a total of 944 high-confidence ciliary proteins and 12 potentially ciliary proteins (red label), resulting in a final set of 956 proteins (Supplementary Table 10) used for the subsequent differential abundance analyses. Using a differential abundance threshold of |log_2_ FC| ≥ 0.5 and adjusted *P*-value < 0.05, we identified 141 ciliary proteins as significantly upregulated and 27 as significantly downregulated at 28 °C compared to 18 °C (Figure 4A, Supplementary Table 11). Most notably, there was a strong and coordinated upregulation of the IFT-machinery: five of six IFT-A subunits and three of the 16 IFT-B subunits significantly increased in abundance (Figure 4B). This proteomic signature mirrors some of the transcript levels (*IFT* 140 of complex A and *IFT* 81 of complex B) elevated at 28 °C (Figure 3A). It should be, however, noted that all other transcript levels belonging to the upregulated IFT proteins (*IFT*s 43, 121, 139, and 144 of the IFT-A complex and *IFT*-B 74 and 80), whose transcripts are not indicated in Figure 3A, are also significantly upregulated in cells grown at 28 °C, but with a |log₂ FC| of less than 1 (Supplementary Table 3).

**Figure 4:**
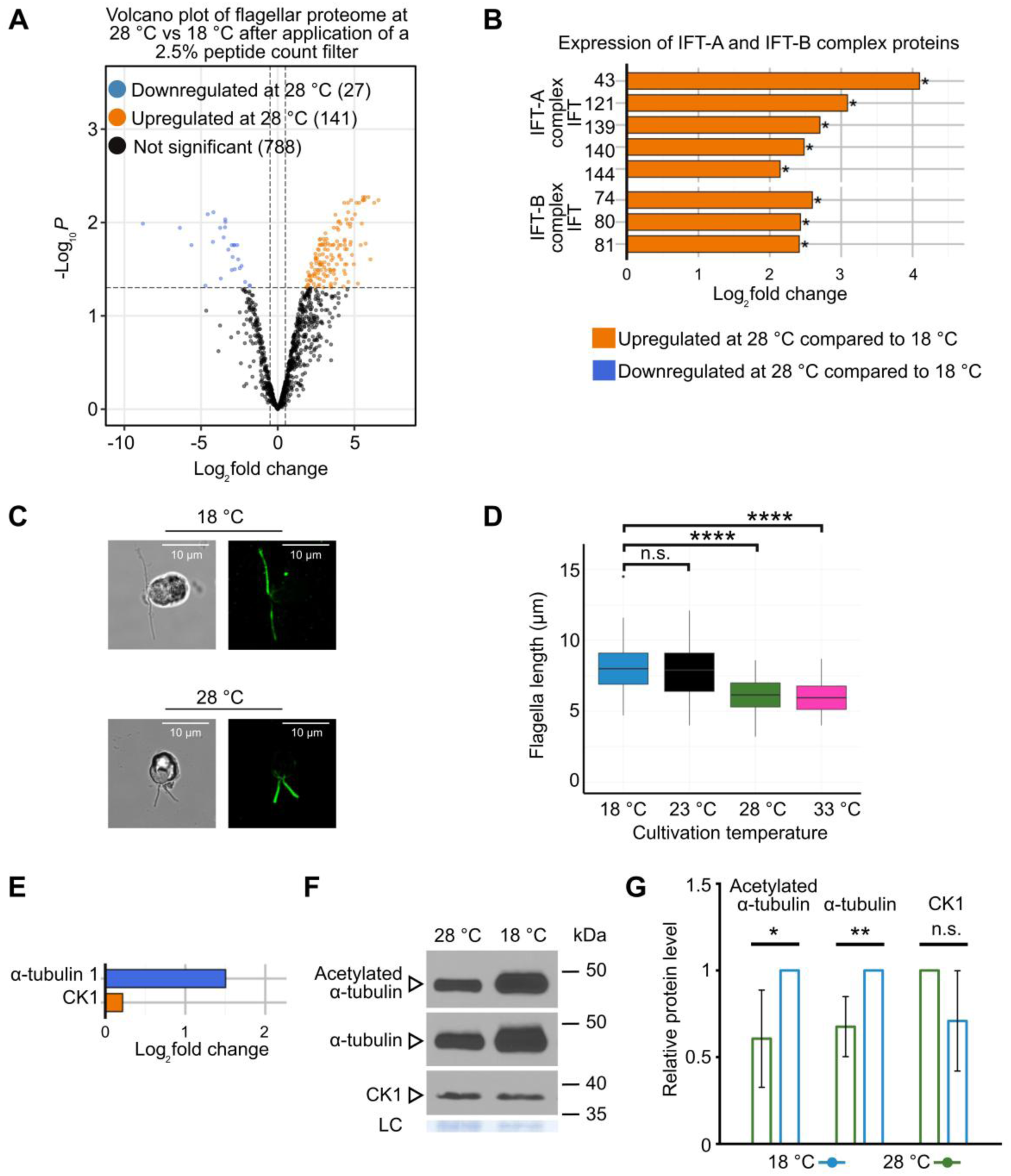
Temperature-dependent remodeling of flagellar proteins alters the length of flagella. **A)** Volcano plot depicting differentially abundant proteins in the flagellar proteome of Chlamydomonas cultivated at 18 °C and 28 °C. Points represent individual proteins. The orange color indicates proteins upregulated at 28 °C, blue color indicates proteins downregulated at 28 °C relative to 18 °C (adjusted *P*-value < 0.05 and |log_2_ FC| ≥0.5), and black represents non-significant changes. Numbers in parentheses denote protein counts per category. Dashed vertical lines indicate ± log_2_ fold change of 0.5; the horizontal dashed line marks the significance threshold. **B)** Barplot showing log_2_ FC in IFT-A and IFT-B complex proteins that were significantly differentially abundant in cells grown at 28 °C compared to 18 °C (adjusted *P*-value < 0.05 and |log_2_ FC| ≥ 0.5). Orange bars indicate upregulation in IFT component abundance, suggesting temperature-dependent modulations in the IFT machinery. Adjusted *P*-values < 0.05 (*), as calculated by FragPipe, are shown. Proteins along with their abbreviations are listed in Supplementary Table 5 and are also indicated in Supplementary Table 11. **C)** Representative immunolocalization microscopy images of Chlamydomonas cells cultivated at either 18 °C or 28 °C. Left panels show brightfield microscopy images of cells and cilia, right panels depict immunofluorescence images of cilia stained with anti-acetylated α-tubulin antibodies (see the Methods section). **D)** Quantification of flagellar length in cells grown at 18 °C, 23 °C, 28 °C, and 33 °C on day 4, measured using immunolocalization images. Each boxplot includes measurements from three biological replicates (50 flagella per replicate; 150 per temperature). Significant differences between temperature treatments were determined by a two-sided *t*-test assuming unequal variance (Welch’s *t*-test) and are marked by asterisks (n.s.: not significant; ****: *P* < 0.0001). **E)** Quantification of α-tubulin and CK1 protein abundance within isolated flagella, shown as a barplot from proteome data. **F)** 0.5 µg (for blots with acetylated α-tubulins and α-tubulins) and 20 µg (for blots with CK1) of NP-40 treated ciliary proteins per lane isolated from cells grown at 18 °C and 28 °C, respectively, were separated by SDS-PAGE and immunoblotted with (i) anti-acetylated α-tubulin, (ii) anti-α-tubulin or (iii) anti-CK1 antibodies as indicated (see Methods). The loading control (LC) of the Coomassie stained membrane used for CK1 detection is shown. **G)** Quantifications of the protein levels of acetylated and non-acetylated α-tubulins as well as CK1 from six biological replicates deriving from five independent datasets are shown. The highest level of each protein was always set to 1. Asterisks represent significant differences as calculated by a two-sided *t*-test assuming unequal variance (Welch’s *t*-test). *P* < 0.05 (*), *P* < 0.01 (**) and not significant (n.s.).

It is known that Chlamydomonas can alter flagella length by activating axonemal disassembly and stimulating IFT trafficking (Marshall and Rosenbaum 2001; Pan and Snell, 2005). To determine whether the observed protein trends translate into a morphological phenotype, we performed immunolocalization of cilia using anti-acetylated α-tubulin antibodies (Figure 4C). On day 4 of growth, flagellar length was the longest in cells cultivated at 18 °C, reaching a mean flagellar length of 8.15 µm at LD6 (see Methods). As the cultivation temperature increased, flagellar length progressively reduced, reaching an average of 6.14 µm at 28 °C and 6.03 µm at 33 °C (Figure 4D, Supplementary Table 12). This reduction in mean length could reflect a highly variable population, with significant numbers of cells possessing very short, regrowing flagella alongside with cells with full-length flagella. To test this alternative hypothesis, we generated frequency distribution histograms of flagellar lengths (Supplementary Figure 9). These data show a unimodal distribution at all temperatures. Thus, the observed phenotype suggests that the entire population acclimates to warmer temperatures by shifting to a progressively shorter mean flagellar length. Comparable results on the reduced length of flagella were observed on day 3 and day 7 of growth at LD6, while levels on day 1 showed no reduced ciliary length (Supplementary Figure 10, Supplementary Table 12).

We also verified whether the reduction in ciliary length correlates with the abundance of a primary ciliary component, α-tubulin. As the distal tip region lacks the standard nine microtubule doublets (Reynolds et al., 2018; Wingfield and Lechtreck, 2018), shorter cilia displaying the complete tip structure would have a reduced relative abundance of α-tubulin compared to full-length cilia.

We observed a small but non-significant decrease in α-tubulin 1 abundance, comparing its level in cells grown at 18 °C versus 28 °C, as measured by LC-ESI-MS/MS (Figure 4E). However, flagellar α-tubulin differs from cell body α-tubulin, as it gets post-translationally modified by acetylation (L’Hernault and Rosenbaum, 1985). To verify the amount of acetylated α-tubulin versus α-tubulin in the flagella, we performed immunoblots with flagellar proteins along with anti-acetylated α-tubulin and anti α-tubulin antibodies, respectively (Figure 4F). As loading control and for comparison, we used antibodies against casein kinase 1 (CK1). LC-ESI-MS/MS measurements show that CK1 is slightly but non-significantly increased in cells grown at 28 °C (Figure 4E). The immunoblot analyses of six biological replicates showed a significant reduction of acetylated α-tubulin as well as of α-tubulin in cells grown at 28 °C (Figure 4F, G). In contrast, CK1 levels were slightly but non-significantly enhanced in cells grown at 28 °C (Figure 4F, G). These data indicate a conserved ciliary tip with a reduced amount of α-tubulin in the shortened cilia, as discussed later.

### Temperature-driven changes in flagellar proteins connected to motility alter algal motility patterns

Further analysis of the flagellar proteome revealed substantial temperature-dependent remodeling of Chlamydomonas’ motility machinery (Figure 5A). A plethora of major motor proteins, including axonemal and cytoplasmic dynein heavy chains, kinesin motor proteins, and dynein light/intermediate chains, exhibited pronounced upregulation at 28 °C, suggesting a heightened demand for active transport and mechanical work within the flagellum at elevated temperatures. Similarly, proteins associated with ciliopathy syndromes, especially the Bardet–Biedl syndrome (BBS) proteins, showed strong upregulation at 28 °C. BBS proteins are connected to IFT trains and influence phototactic behavior (Liu and Lechtreck, 2018). In contrast, ankyrin repeat proteins (FAP164 and FAP208), which are critical for maintaining axonemal stability and regulating ciliary and flagellar architecture, displayed reduced abundance. Moreover, downregulation of two zinc chaperones (ZNG1, ZNG2) was observed in cells grown at 28 °C. A similar trend was detected in the GO enrichment analyses, since protein folding and protein refolding were enriched GO terms, with many transcripts showing downregulation at 28 °C (Figure 1G). Such chaperones are important for maintaining protein homeostasis and might play a role in flagellar assembly under changing environmental conditions, also at lower temperature. Beyond, we identified signaling proteins including kinases and a PAS domain protein. One of them is the Neural-Wiskott Aldrich syndrome protein bearing a kinase domain (Phytozome 14) that is involved in cilia biogenesis in mice (Jain et al., 2014). Another one, the aurora-like kinase, CALK, is linked to flagellar length regulation through phosphorylation (Luo et al., 2011). Taken together, the proteome from cells grown at 28 °C reveals more dynamic flagella with heightened intraflagellar transport and increased abundance of axonemal motors.

**Figure 5:**
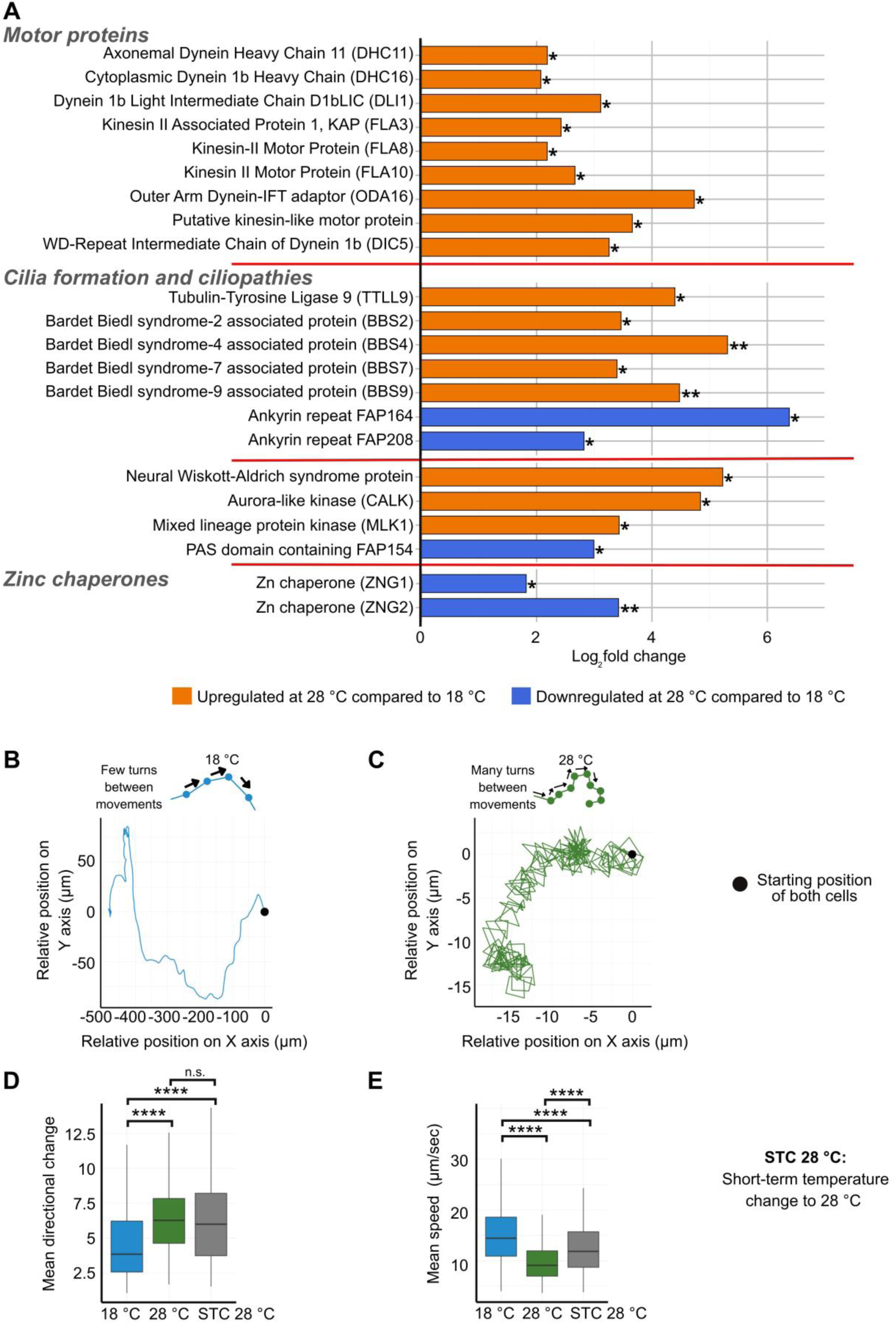
Remodelling of ciliary proteins and changes in swimming speed and algal cell rotations at different ambient temperatures. **A)** Log_2_ fold change of selected flagellar proteins from multiple functional groups in Chlamydomonas compared between cultivation temperatures of 18 °C and 28 °C. Motor proteins (e.g., dyneins, kinesins), ciliopathy-related proteins (BBS proteins, ankyrins), kinases or zinc chaperones display significant upregulation or downregulation, as indicated by colored bars. Orange bars denote proteins upregulated at 28 °C relative to 18 °C, while blue bars indicate downregulation. Statistically significant proteins are marked by asterisks. Adjusted *P*-values < 0.05 (*) and < 0.01 (**) are shown, as calculated by FragPipe. Proteins along with their abbreviations are listed in Supplementary Table 5 and are also indicated in Supplementary Table 11. **B, C)** Motility paths tracked for a representative Chlamydomonas cell cultivated either at 18 °C (B) or at 28 °C (C). The first 60 s of the movement are depicted. Please note that the x-axes in B and C show different scales. Cells at 18 °C typically exhibit fewer turns between movements, resulting in relatively straight trajectories. In contrast, cells at 28 °C display frequent directional changes, producing more curved or meandering movement paths compared to 18 °C. **D)** Quantification of mean directional change (degrees) between frames (i.e. at 200 ms intervals) for cells grown four days at 18 °C and 28 °C, respectively, or grown at 18 °C for four days and being then exposed for 15 min to 28 °C (STC). Data reflects a minimum of 150 individual tracks per temperature per biological replicate, with three biological replicates in total for 18 °C, two biological replicates for 28 °C, and two biological replicates for STC. The mean of all biological replicates is shown. The two and three, respectively, independent biological replicates are shown in Supplementary Figure 11. **E)** Mean swimming speed (µm/sec) as calculated from tracked cell movements for cells grown four days at 18 °C and 28 °C, respectively, or grown at 18 °C for four days and being then exposed for 15 min to 28 °C. Data reflects a minimum of 150 individual tracks per temperature per biological replicate, with three biological replicates in total for 18 °C, two biological replicates for 28 °C and two biological replicates for STC. The mean of all biological replicates is shown. The two and three, respectively, independent biological replicates are shown in Supplementary Figure 11. The speed is calculated by computing the mean swimming speed along each trajectory (in the x-y plane) then averaging this value over trajectories. **D, E)**, Statistical significance determined by a two-sided *t*-test assuming unequal variance (Welch’s *t*-test) is indicated by asterisks; *P* < 0.0001 (****), n.s. not significant.

We also sought to investigate the possible behavioral consequences of these molecular changes in flagellar-related proteins, by taking a close look at cell motility. For this purpose, we examined the motility of Chlamydomonas cells using microscopy in a temperature-controlled chamber, combined with the ImageJ plugin TrackMate (see Methods), which quantifies various parameters associated with cell trajectories, including average speed and turning angle, the latter indicating to what extent cells change their direction between frames. We used the same cell densities for all samples (see Methods and Discussion). Representative trajectories for individual cells grown at each temperature illustrate strong shifts in motility patterns. Many cells grown at 18 °C followed relatively straight paths, with infrequent directional changes (Figure 5B, Video 1). In contrast, many cells grown at 28 °C exhibited more constrained paths characterized by frequent apparent abrupt directional changes (Figure 5C, Video 2). Figure 5B and C show the trajectories of the individual cells for 60 s; the videos show six examples of tracked cells for a total of 57 s. Quantitative analysis of over 150 tracked cells per biological replicate experiment confirmed these observations. It should be noted that the basic swimming behavior of Chlamydomonas involves helical movement with breaststrokes when moving forward (Crockett, 2021). This ensures that the eyespot situated at the nearby *cis*-flagellum perceives light from all directions. The directional change between frames, averaged over a trajectory, was significantly higher in cells grown at 28 °C compared to 18 °C (Figure 5D, left and medium panels). We also calculated the mean swimming speed along each trajectory. Averaging over trajectories, this was significantly reduced in cells cultivated at 28 °C compared to 18 °C (Figure 5E, left and medium panels), consistent with the assumption that frequent course corrections lead to slower cell speed. We investigated whether the observed motility differences were a consequence of the four-day long acclimation or were rather an immediate response to the temperature shift. To find this out, we transferred cells that had been grown at 18 °C for four days to a short-time temperature change (STC) of only 15 min to 28 °C and analyzed them directly for the mean directional change (Figure 5D, right panel) and mean speed (Figure 5E, right panel) for 15 min on the temperature-controlled microscope slide. Our data clearly show that the STC to 28 °C is enough to alter the mean directional change pattern to the one observed in cells grown for four days at 28 °C (Figure 5D, right panel). In contrast, mean speed after the STC was significantly different to that observed for cells grown for four days at 28 °C (Figure 5E, right panel), but the cells were on average still slower than cells that had been grown for four days at 18 °C. Thus, speed is determined to some extent by acclimation, while the turning pattern depends mainly on the current temperature.

### Temperature-driven changes in the Chlamydomonas secretome and consequences for mating efficiency

Transcriptomic analyses revealed pronounced downregulation of transcripts encoding ECM components at higher temperatures, specifically the pherophorins and MMPs, that are all linked to the secretome (Luxmi et al., 2018). Thus, in the next step, we analyzed whether the Chlamydomonas secretome also changed with cultivation temperature, using label-free quantitative proteomics (see Methods). Mass-spectrometric profiling of Chlamydomonas proteins in supernatants (see Methods) of cells cultured at 18 °C and 28 °C was performed with a 2.5% peptide count filter, as applied for the ciliary proteome (see Methods). With a maximum identification of 110 unique peptides per protein in the secretome, a protein required at least three unique peptides per protein to qualify. We identified a total of 1,764 proteins with at least three identified peptides (Supplementary Table 13). We analyzed these 1,764 proteins using DeepLoc to predict extracellular proteins. Of the 1,764 proteins detected, 455 were predicted by the program as extracellular proteins and constituted our final dataset of secreted proteins (Supplementary Table 14). Using thresholds of |log_2_ FC| ≥ 0.5 and adjusted *P*-value < 0.05, we found that 286 proteins were significantly upregulated and 39 were significantly downregulated at 28 °C relative to 18 °C (Figure 6A, Supplementary Table 15).

**Figure 6:**
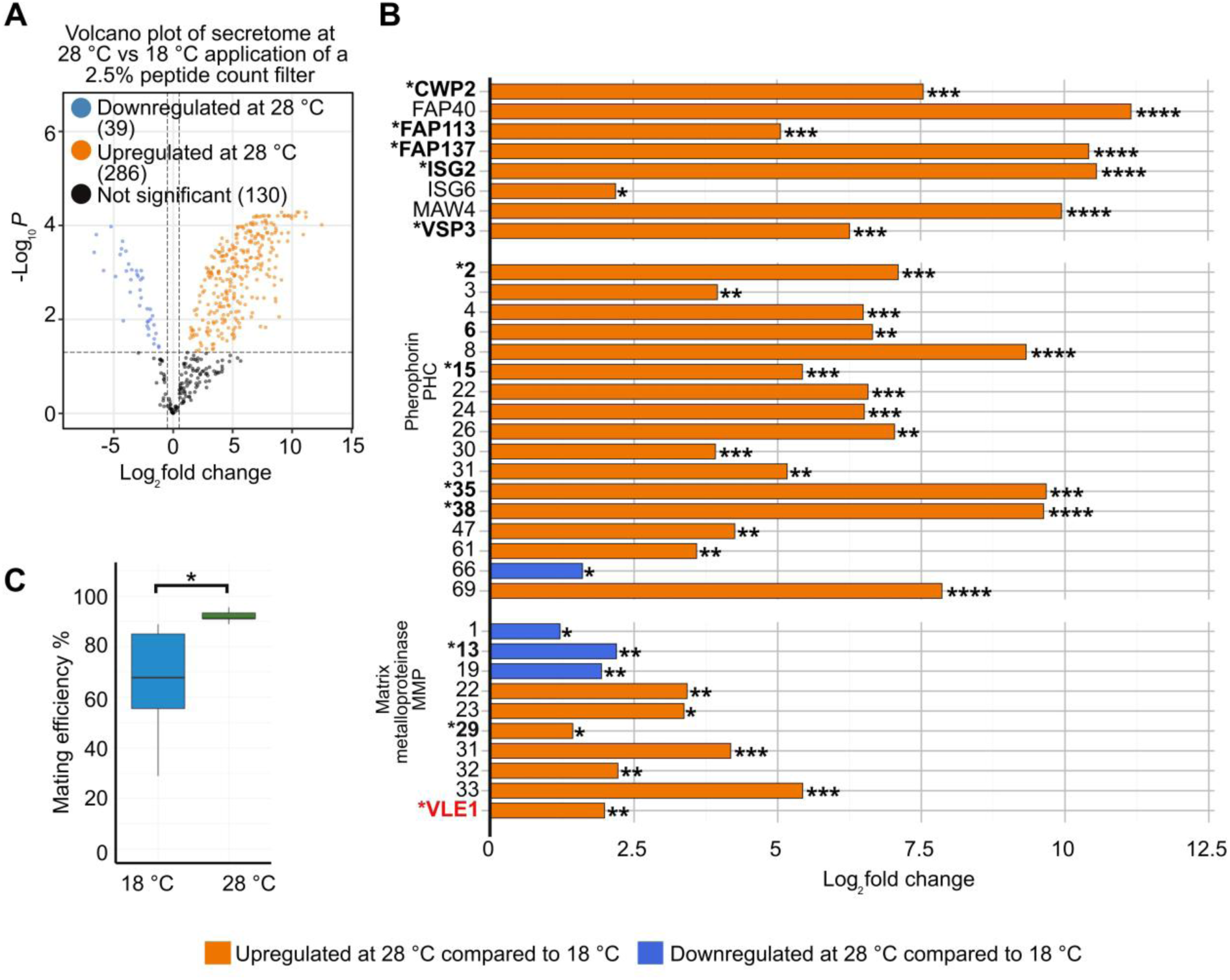
Temperature effects on the secretome, extracellular proteins, and mating efficiency in Chlamydomonas. **A)** Volcano plot depicting differentially abundant proteins in the Chlamydomonas secretome cultivated under 18 °C and 28 °C. Each point represents a unique protein; orange and blue indicate significant up- or downregulation. Axes indicate log_2_ fold change versus –log_10_ (adjusted *P*-value); numbers in parentheses denote protein counts per category. **B)** Barplot showing the log_2_ FC in abundance of key extracellular proteins as predicted by DeepLoc in the secretome. Notably, the protein VLE1 was detected among soluble mating secretory proteins in previous studies (Luxmi et al., 2018) but was not predicted as extracellular by DeepLoc. Other proteins detected in the soluble mating secretome are highlighted with an asterix and in bold. Adjusted *P*-values < 0.05 (*), < 0.01 (**), < 0.001 (***), and < 0.0001 (****) are shown, as calculated by FragPipe. Proteins along with their abbreviations are listed in Supplementary Table 5 and are also indicated in Supplementary Table 15. **C)** Quantification of mating efficiency across temperature or experimental conditions. Data represents three biological replicates. The blots show the proportion of successfully fused mating pairs, with mean values marked. Significant differences in mating efficiency between groups were determined using a two-sided *t*-test assuming unequal variance (Welch’s *t*-test); *P*-values are indicated (*: *P* < 0.05). A second independent dataset with three biological replicates is shown in Supplementary Figure 12.

While many proteins of the cilia coincide in their transcript and protein abundances, the secretome responds in the opposite direction to the transcriptome. Many transcripts encoding ECM proteins and other secretory proteins are downregulated (Figure 3B) or unchanged, but a large number of secreted proteins, including HRGPs, pherophorins and MMPs are strongly upregulated in cells grown at 28 °C relative to 18 °C (Figure 6B). We also compared the ambient temperature-altered secretomes against the published Chlamydomonas mating secretome, measured at 22 °C (Luxmi et al., 2018), and identified interesting overlaps between the two datasets. The ambient temperature-altered secretome contains numerous proteins with established roles in mating, including multiple pherophorins, gametolysins, and subtilisin-like proteases (these are all marked with a * in front of the name in Figure 6B). The differential regulation of VLE1, a key subtilisin-like protease that is also released during gametogenesis, was particularly relevant, as discussed later. The temperature-dependent regulation of gametogenesis-related proteins could prime cells for developmental programs that are advantageous under warmer conditions, potentially accelerating the onset of mating competence.

To test this hypothesis functionally, we performed mating experiments using Chlamydomonas strains CC-124 (mt^-^) and CC-125 (mt^+^), routinely used for mating assays in our laboratory (Müller et al., 2017). Strains were cultivated under nitrogen-limited conditions at different ambient temperatures (see Methods). Mating efficiency assays revealed that higher cultivation temperatures significantly increase gamete fusion success, with cultures pre-conditioned at 28 °C showing enhanced mating competency compared to those grown at 18 °C (Figure 6C).

## DISCUSSION

We used transcriptome analysis to uncover gene expression changes in the biflagellate model microalga Chlamydomonas acclimated to different ambient temperatures. We complemented these data with proteomic and behavioral analyses that highlighted structural changes in cilia impacting the swimming speed. In contrast, swimming trajectories are affected even by short-term temperature changes. We observed shifts in algal-bacterial interactions, photoreceptor regulation, carbon dioxide consumption and marked changes in the secretome, connected to enhanced mating behavior, at higher temperatures. The temperature ranges investigated included 18 °C, 23 °C, 28 °C, and 33 °C for transcriptomic analyses, while our proteomic analyses focused on 18 °C and 28 °C. These conditions differed from those in previous system-level studies, which primarily addressed moderate or acute heat stress at 35 °C and 40 °C, compared to normal (25 °C) and heat-stressed growth (Zhang et al., 2022; Zhang et al., 2024), or examined cold stress below 15 °C (Valledor et al., 2013). Anticipating that climate change will both shift the typical range of temperatures and increase extreme stress situations, it is central to understand the algal cellular responses over both stress-inducing and ambient temperature ranges. In their natural habitat of flooded rice fields, Chlamydomonas has historically been exposed to temperatures ranging from 15 °C to 30 °C (Chin et al., 1999), as well as to acetate as major carbon source (Hori et al., 2007). A range of temperatures extending 3 °C above the natural limits experienced by Chlamydomonas were tested here, allowing us to investigate the effects of rather moderate warming without exposing the algae to heat stress. We also simulated the natural environment by incubating the algal cells in acetate-containing medium (Harris, 1989). We checked acetate levels in the culture medium on day 4 when the cells were harvested for the transcriptome and proteome studies at the used temperatures of 18 °C and 28 °C (Figure 3F). The levels were reduced compared to day 1, but there was still acetate present in the cultures grown at both temperatures. In our experiments, we aimed to study the effects of the different temperatures on different parameters like acetate consumption in relation to CO_2_ uptake, rather than creating laboratory conditions that are suited for optimal growth over the entire experimental period.

Transcriptome profiling revealed that ambient temperature drastically remodels the algal transcriptome. The scale of this change was extensive: compared to cells grown at 18 °C, cultivation at 28 °C resulted in over 5000 differentially abundant transcripts, while cultivation at 33 °C affected more than 6000 transcripts. In both comparisons, this represents roughly one-third of all transcripts in the genome (vs 6.1, Craig et al., 2023).

As stated earlier, TRP channels and photoreceptors can sense temperature. While missing in land plants, animals as well as some algae have a high number of TRP channels (Verret et al., 2010). 18 of the 29 annotated TRP channel transcripts based on vs 6.1 of the Chlamydomonas genome (Hou et al., 2023) display significant differential abundance at 28 °C or 33 °C compared to 18 °C with a |log₂ FC| ≥ 1. Especially interesting findings include the changes in transcript abundances encoding the TRP channels 5, 11, and 22 that are involved in the antagonistic interaction of the bacterium *P. protegens* against Chlamydomonas (Hou et al., 2023). These three channels group close to the vanilloid family (TRPV) tree that includes thermosensitive and calcium-permeable members in mammals, also playing a role in autoimmune diseases (Xiao et al., 2025). Among them, the *TRP5* transcript is most strongly downregulated in cells grown at 33 °C, but also shows a significant decrease at 28 °C. *TRP5* transcript abundance also decreases, in conjunction with a thickening of the algal cell wall, when algal cells are transferred to 3-dimensional (3-D) conditions in the presence of acetate to simulate nature-like conditions in rice fields (Vuong et al., 2025). Since algal fitness against antagonistic bacteria is enhanced under 3-D conditions, we analyzed whether ambient temperature changes would similarly influence algal-bacterial antagonism. We found that higher ambient temperatures indeed enhance algal fitness (Figure 2B - D, Supplementary Figure 5). This improved resilience is likely due to an interplay between internal algal molecular characteristics and extrinsic bacterial growth dynamics. Internal algal molecular characteristics likely confer protection through the strong temperature-dependent downregulation of *TRP5* and *TRP11* transcripts. Based on our prior studies (Hou et al., 2023), we hypothesize that the decrease of these two channel transcripts, relative to the slight increase of *TRP22*, reduces the deflagellation rate upon exposure to the bacterial toxin orfamide A. Additionally, internal defense may be supported by a strengthened physical barrier, indicated by the increased abundance of ECM-related proteins found in the secretome at higher temperatures (Figure 6B). On the other hand, extrinsic bacterial growth dynamics plays a critical role in the interaction’s outcome. While *P. protegens* grows rapidly at higher temperatures, it exhibits a sharp decline in cell density after reaching an early maximum. At 28 °C and 33 °C, bacterial density peaks around day 1 and subsequently crashes, resulting in low bacterial pressure by the time algae begin to recover (Figure 2E). In contrast, at 18 °C, bacterial cell density peaks later and declines much more slowly, maintaining a higher antagonistic pressure on the algae throughout the cultivation period (Figure 2E). The knowledge about this temperature-dependent algal-bacterial antagonism is likely to be transferable to other ciliated algae that are sensitive to orfamide A (Aiyar et al, 2017), including the freshwater species *Haematococcus pluvialis* and *Gonium pectorale* as well as the marine *Chlamydomonas* sp., whereby the latter two are known to be part of algal blooms (Aiyar et al, 2017; Saggiomo et al., 2021).

20 known/predicted photoreceptors are encoded in the genome of Chlamydomonas (Greiner et al., 2017; Sharma et al., 2025). In land plants, it was shown that photoreceptors sense temperature changes (reviewed in Sharma et al., 2024). Our data (Figure 2G) emphasize that at least four different algal photoreceptors can sense ambient temperature variations at the protein level. For two photoreceptors, CHR1 and CRY-DASH1, changes in transcript abundance were positively verified at the protein level. While CHR1 is being upregulated at higher temperatures with an optimum at 28 °C, the CRY-DASH1 level is highest at 18 °C (Figure 2G). CHR1, a microbial channelrhodopsin, emerged as a major tool in modern biological research and was one of the first algal photoreceptors used for optogenetic approaches (Deisseroth and Hegemann, 2017; Christie and Zurbriggen, 2021). In the algal cell, CHR1 is involved in phototactic behavior and is situated in a specialized part of the plasma membrane directly adjacent to the eyespot (Schmidt et al., 2006). The other photoreceptor, CRY-DASH1, helps balancing the photosynthetic machinery by acting as a negative regulator that prevents hyperstacking of thylakoid membranes (Rredhi et al., 2021). In addition, it is involved as a positive regulator in anoxic metabolism (Rredhi et al., 2024). We also found two further photoreceptors that sense temperature; they seem to be regulated at the posttranscriptional/translational level. These are aCRY, a blue- and red-light perceiving receptor (Beel et al., 2012) and CRY-DASH2 whose function is still unknown. While aCRY is upregulated at 18 °C like CRY-DASH1, CRY-DASH2 is upregulated at 28 °C and 33 °C (Figure 2G). The inverse accumulation of the two CRY-DASH proteins across the physiological temperature range are reminiscent of the antagonistic regulatory mechanisms that underlie temperature compensation in circadian clocks (Mittag et al., 2005). To achieve compensation, such biological oscillators may use an antagonistic pair of processes to counterbalance thermal effects and ensure reliable timekeeping. Double knockout mutants of the *CRY-DASH* genes may help to find out in the future whether the two proteins may be involved in this process.

Studies of transcripts controlling metabolic pathways highlighted that both lipid biosynthesis, specifically betaine lipid metabolism, and proteins involved in photosynthesis and chlorophyll biosynthesis are upregulated at 18 °C. Betaine lipids such as DGTS may facilitate this adaptation by serving as carriers for polyunsaturated fatty acids (PUFAs) and maintain membrane-bilayer homeostasis. It was surprising that this potential mechanism to increase membrane fluidity is triggered at temperatures as high as 18 °C, and not only under severe cold-stress conditions.

CO_2_ consumption and production measurements also revealed interesting new aspects. When acetate is available, algal cells are known to grow mixotrophically (Harris 1989). The cells grown at 18 °C clearly exhibited photoautotrophic growth during the four-day measurement period (Figure 3E), as evidenced by the small but consistent net CO_2_ consumption (negative CTRs) during light phases being most obvious on day 3, and low acetate consumption by day 4. In contrast, cells grown at 28 °C appeared to prioritize heterotrophic growth at first. During the first three days, these cells showed no negative CTRs even during light phases, although their cell number is higher compared to the one at 18 °C. However, on day 4, a switch to net photoautotrophy (negative CTRs) occurred, after the CO_2_ content was strongly increased in the night phase of day 3. This coincided with the reduction of acetic acid (Figure 3F). These findings suggest that elevated temperatures potentially combined with organic carbon availability can delay the onset of net photosynthesis. In warming, nutrient-enriched waters, this metabolic shift could have severe consequences for the ecosystem. If mixotrophic algae spend more energy on respiring rather than photosynthesizing, the ecosystem loses a critical period of carbon dioxide sequestration and oxygen production. For all O_2_ consumers, including humans, this reduction in algal oxygen generation would be a significant disadvantage. Alternatively, this metabolic plasticity may have significant implications for algal-based wastewater treatment. The results suggest that maintaining elevated cultivation temperatures (approximately 28 °C) drives the algae to focus on the uptake of organic carbon rather than CO_2_ fixation.

As many transcripts involved in the ciliary structure and in the cellular secretory processes were modified at different temperatures, we extended our studies to include the ciliary and secretory sub-proteomes. Proteomic analysis of the NP-40 treated ciliary proteins isolated from cells grown at 18 °C and 28 °C, respectively, revealed a strong increase of IFT-A and IFT-B complex proteins, which are crucial for intraflagellar transport from the base of the cilia up to the tip (anterograde IFT train) and from the tip down to the basal body (retrograde IFT train) (Lacey et al., 2024). The increase in IFTs observed at the transcriptome and protein levels (Figures 3A and 4B, Supplementary Table 3) coincides with the upregulation of many motor proteins such as kinesin II motor proteins FLA8, FLA10 and KAP1 (FLA3) and several dyneins (Figure 5A) at higher temperature while the post-translationally modified acetylated α-tubulin as well as α-tubulin are downregulated under these conditions (Figure 4E-G).

The proportional reduction in α-tubulin abundance in shortened cilia of cells grown at 28 °C can be explained inter alia by the structural composition of the ciliary tip region. The ciliary axoneme has the typical 9+2 structure with nine doublet microtubules and two central pair singlets (Pazour et al., 2005). The microtubules consist mainly of α- and ß-tubulins. The distal tips of regrowing and mature cilia lack the standard nine microtubule doublets (Reynolds et al., 2018; Wingfield and Lechtreck, 2018). Instead, this region contains only the central pair singlets along with nine extended microtubule singlets. At the top of the tip, only the central pair is still present ending in the flagellar tip complex. Shortened cilia that maintain the tip architecture will therefore contain proportionally less total tubulin (Figure 4E-G).

The remodeling of many ciliary protein abundances in cells grown at 28 °C correlates with shorter cilia observed at this temperature in the middle of the day (LD6, Figure 4C and D). Of special interest is CALK that determines ciliary length via its phosphorylation status (Luo et al., 2011). CALK is mainly present in the algal cell body but is also found in cilia. It is upregulated at 28 °C. The control of flagellar length in Chlamydomonas is also related to the diurnal cycle (Tuxhorn et al., 1998) and is linked with IFT particles as well as axonemal disassembly. This process has been discussed in various models (Marshall & Rosenbaum, 2001; Pan and Snell, 2005; Wemmer et al., 2020; Patel et al., 2024). In the future, it will be interesting to study whether IFT particles may move differentially at different temperatures. Moreover, four proteins of the BBS complex (BBS-2, −4, −7, and −9) related to ciliopathies are increased in abundance in cells grown at 28 °C (Figure 5A). Previous studies found that a purified BBS-like complex containing BBS1, −4, −5, −7, and −8 associate with a subset of IFT proteins (Lechtreck et al., 2009). Three of these IFT-associated BBS proteins are also temperature-regulated. Interestingly, some Chlamydomonas BBSome mutants (*bbs1*, *bbs4*, *bbs7*), can assemble full-length, motile flagella but become completely non-phototactic, due to an unregulated accumulation of critical signaling proteins (Lechtreck et al., 2009). These results may thus also connect with the observed modified swimming behavior at 18 °C and 28 °C (Figure 5 B-E). Behavioral changes in swimming trajectories have been found before under cold and heat stress conditions and depend further on the acclimation duration (Meier et al., 2022). In addition to temperature and acclimation, population density plays an important role in the determination of swimming speed. The speed increases with higher cell densities due to the secretion of a yet unknown low-molecular weight compound (Folcik et al., 2020). Here, we used the same cell density for all measurements and kept it relatively low (5 × 10^5^ cells mL^-1^) for comparative studies. This density was selected due to the limited growth of cultures at 18 °C by day 4 after inoculation (Figure 1A, Methods). The calculated speeds are in the range of priorly observed speeds of Chlamydomonas cells based on similar cell numbers (Folcik et al., 2020). Using a specialized temperature-controlled microscopy setup (see Methods), we revealed that even ambient changes in temperature result in severe alterations in swimming speed as well as in the frequency of turns performed by the algal cells.

The temperature-dependent differences in swimming speed and directional changes may be due to acclimation processes or may be a direct consequence of the temperature. Short-term temperature change experiments suggest that directional changes are indeed a direct consequence of the temperature (Figure 5D). In contrast, the swimming speed depends at least to some degree on acclimation (Figure 5E). As the differences in flagellar length at the different temperatures are not obvious on day 1 of growth (Supplementary Figure 10), they seem not to be related to the observed directional changes on short term. They may, however, be involved in swimming speed. Cell length is already altered on day 1 (Supplementary Figure 1), which could play a role in these processes. It is impressive how quickly ambient temperature changes control behavioral and morphological processes in Chlamydomonas. We assume that other ciliated algae may be also prone to such changes and that climate change will influence algal behavior and thus ecological systems on a wide range.

The analysis of the secreted proteome revealed a striking temperature-dependent reorganization in Chlamydomonas characterized by the substantial upregulation of MMPs, the vegetative lytic enzyme 1, VLE1, and a diverse array of pherophorins at 28 °C, creating a molecular environment primed for extensive ECM modification. MMPs belong to a family of zinc-dependent proteinases and can cleave diverse substrates including components of the ECM (Hey and Linder, 2024). One of the first MMPs of Chlamydomonas that was characterized represents the gamete lytic enzyme (Kinoshita et al., 1992), but several MMPs and other proteolytic enzymes such as VLE1 have been also found in the mating secretome performed at 22 °C (Luxmi et al., 2018). Pherophorins are a major group of extracellular hydroxyproline-rich proteins in Chlamydomonas (Barolo et al., 2022) and are also important in *V. carteri* in ECM biosynthesis (von der Heyde and Hallmann, 2022). 16 pherophorins are upregulated at higher temperature as well as eight further hydroxyproline-rich proteins, including CWP2, FAPs 40, 113 and 137, ISGs 2 and 6, MAW4 and VSP3. We analyzed whether the increased secretion may be correlated with a higher cell lysis rate at 28 °C by staining cells with Evans Blue. If cells are not intact, the dye can penetrate them. There was a minor degree of staining with no statistically significant difference in cells grown at 18 °C or 28 °C (Supplementary Figure 13), highlighting that the increase in ECM is not related to cell lysis.

Several of the secretory proteins (marked by an asterisk in Figure 6B) overlap with proteins identified in the Chlamydomonas mating secretome study (Luxmi et al., 2018). Sexual algal reproduction was shown earlier to depend on the secreted protein profile (Snell et al., 1989). Our results highlight that cultivation at elevated ambient temperatures significantly enhances the secretion of proteins required for mating. Thus, we examined the mating efficiency of plus and minus gametes at different temperatures and found that gametogenesis increases as temperature rises (Figure 6C). These data suggest an influence of the ambient temperature changes on the sexual life cycle of Chlamydomonas. Such changes seem not limited to Chlamydomonas. Early on, it was shown that the chlorophyte *Protosiphon botryoides* needs less time for gamete formation when the temperature is increased from 18.5 °C to 28 °C (Maher, 1947). Also, *Chlamydomonas chlamydogama* has a higher percentage of zygospore formation at increased ambient temperatures of 22 °C, 26 °C, 30 °C, and 35 °C (Trainor, 1960). The higher temperatures may act as an anticipatory signal for deteriorating environmental conditions in summer times. Notably, it has been shown for Chlamydomonas that the germination efficiency of zygospores is controlled by the photoperiod; it is enhanced under long day light-dark cycles representing summer, when temperatures are in general higher (reviewed in Mittag et al., 2005).

The ciliary and secretome proteomes identified more proteins (Supplementary Tables 9 and 13) than are present (i) in the used ciliary databases along with few candidates of unknown function (Supplementary Table 10) and (ii) predicted to be localized extracellularly by DeepLoc (Supplementary Table 14), respectively. We are aware that these proteins may be contaminants.

To strengthen our proteome data, we applied a 2.5% peptide filter threshold to both datasets. In case of the ciliary proteome, we could include 210 ciliary proteins positively verified against further existing ciliary databases, as ciliary proteins are well conserved (reviewed in Li et al., 2025), or by current literature (Supplementary Table 9). In contrast, the secretome posed a greater challenge for comparative analysis, as secreted proteins vary significantly in composition and quantity between organisms, even among plants (Alexandersson et al., 2013). Consequently, we primarily relied on DeepLoc predictions for localization. This prediction, however, can miss known secreted enzymes. For example, VLE1, a well-characterized secreted Chlamydomonas protein (Luxmi et al., 2018), was not predicted by DeepLoc to localize to the extracellular region. Based on literature, VLE1 was included in our list. We did not include secretory proteins from other organisms. Still, we would like to mention that proteins can perform distinct cellular functions in different compartments, also known as protein moonlighting. This is particularly known among extracellular proteins such as mammalian heat shock proteins (HSPs) (Jeffrey, 2017). In the Chlamydomonas secretome, several HSPs were identified with high peptide counts, yet only one passed DeepLoc’s extracellular prediction criteria.

In some cases, the transcriptome changes agree with changes found at the protein level, but in other cases, like with some photoreceptors or especially with the secretome, correlations are scarce or even contradictory. Thus, post-transcriptional and translational control mechanisms along with RBPs are most likely involved. The transcriptome studies have shown that transcripts encoding a variety of RBPs are altered (Supplementary Figure 6). Former studies of RBPs selected at different ambient temperatures had revealed that the C3 subunit of RBP CHLAMY1 (CRB3) is downregulated at the protein level in cells grown at 28 °C compared to 18 °C (Li et al., 2018), which agrees with its transcript level (Supplementary Figure 6). Notably, the C3 subunit confers thermal acclimation to Chlamydomonas as shown by mutant lines (Li et al., 2018).

Our present studies highlight that acclimation to different ambient temperatures and even short-term temperature changes represent an important factor in the regulation of a variety of biological processes in Chlamydomonas. Some of the components discussed here, such as TRP channels and photoreceptors are likely to be part of the thermal acclimation pathway involved in sensing ambient temperatures. As some of these receptors are present in other algae, land plants or even mammals, the results can be transferred broadly. Components like the C3 subunit may mediate sensed information. Homologues of C3 like the rat CUG-binding protein2 have been also found in mammals (Zhao et al., 2004). Finally, our study identified components like IFT or ECM proteins that are involved in executing cellular responses like the shortening of the cilia or the enhancement of protein secretion. These events then may induce behavioral changes such as alterations of swimming speed, mating ability or bacterial antagonism. Collectively, our findings demonstrate that ambient temperature changes well outside the typical stress range influence key properties of the organism and demand further focused studies in the future.

## METHODS

### Strains of *Chlamydomonas reinhardtii* (thereafter Chlamydomonas) and *Pseudomonas protegens* Pf-5 used and quantification of growth

Chlamydomonas wild-type strain SAG 73.72 (mt^+^) was obtained from the Culture Collection of Algae at Göttingen University and used for all experiments except for the mating assay. Strain SAG 73.72 is an isolate of Tsubo’s strain C8 that is close to 21 gr from Sager (Harris, 1989). Strain 21 gr served as a crossing partner for generating strain CC-4532, whose genome has been sequenced (Craig et al., 2023).

Cells were cultivated in Tris-Acetate-Phosphate (TAP) medium (Harris, 1989) under a 12 h : 12 h light:dark (LD) cycle using a white light spectrum (Osram L36W/840, lumilux, cool white) at a light intensity of 50 μmol m^-2^ s^-1^ and stirring (100 rpm). Growth experiments were performed under four cultivation temperatures,18 °C, 23 °C, 28 °C, and 33 °C. To initiate precultures, cells were transferred from TAP agar plates (Harris 1989) into 15 mL of liquid TAP medium in sterile NUNC flasks. These were incubated at the designated temperatures with orbital shaking at 100 rpm under the LD cycle for 4 days. On day 4, main cultures were prepared by measuring the cell density of each preculture and adjusting the starting concentration to 1 × 10^5^ cells mL^-1^. This step ensured uniform inoculum density across all temperature conditions. Culture densities were quantified daily by sampling 180 μL of sample and counting cells on an improved Neubauer chamber (no. T729.1; Carl Roth, Karlsruhe, Germany). Visual documentation of chlorophyll content (as a proxy for pigmentation and viability) was also captured every second day by photographing culture flasks. Cell samples for the different experiments were always taken in the middle of the day, at LD6, unless otherwise indicated.

To determine bacterial cell densities in the bacterial cultures, single-use cuvettes were filled with 1200 µL of LB medium for *P. protegens*. Subsequently, 50 µL of 2-day-old *P. protegens* precultures were added to the cuvettes. Using cuvettes containing only media as a blank, the optical density at 600 nm (OD_600_) was used to measure culture density with a spectrophotometer. A calibration formula derived from the linear correlation between OD_600_ measurements and bacterial cell densities was then applied to calculate cell densities of the bacterial precultures (Hotter et al., 2021). Bacterial cultures were standardized to 2.5 × 10^7^ cells mL^-1^ in TAP medium. Cells were grown at the indicated temperature for eight days in a microplate reader (Infinite M Nano+”, Tecan). OD_600_ was recorded every 30 min. TAP medium was used as the blank.

### Statistics

Statistical significance was determined using a two-sided *t*-test assuming unequal variance (Welch’s *t*-test). To account for multiple comparisons, *P*-values were adjusted using the Benjamini-Hochberg (BH) procedure where applicable (Figures 1A, B, 2A, C, D, F, 3A, B, 4B, D, E, 5A, 6B and Supplementary Figures 1A,B, C, D, 4, 5A, b, 6, 10). Error bars in all figures indicate the standard deviation (SD).

### Determination of cell length and estimation of cell areas

Cell lengths were quantified on days 1, 4 and day 7 of growth (Figure 1A). For this purpose, 2 mL aliquots from the main Chlamydomonas cultures were harvested and centrifuged at 500 × *g* for 10 min. The algal cells were fixed with Lugol’s solution and further treated as described previously (Vuong et al., 2025). Brightfield microscopy images along with a Zeiss Axiophot were taken to measure multiple cells with ImageJ (version 1.53t, Schneider et al., 2012) taking the vertical length of the cells into consideration.

In addition to the vertical cell length, cell area estimations were performed on day 1 and 4 of growth. To account for cellular boundaries, cell areas were determined. For this purpose, each cell was meticulously traced by setting corner points using the Polygon Selection tool, available on ImageJ (version 1.53t, Schneider et al., 2012). This procedure provides a rough representation of the cell areas.

### Transcriptome sequencing and data analysis

Total RNA was isolated from cells grown in TAP media under four different ambient temperatures, with three biological replicates per condition. Total RNA was extracted using the Qiagen Plant RNeasy Mini kit (Qiagen, Germany) according to the manufacturer’s instructions. RNA concentration and purity were measured using a NanoDrop ND-1000 UV/Vis spectrophotometer (Thermo Fisher Scientific, USA). Library preparation and sequencing were performed by Eurofins Genomics (Ebersberg, Germany). For each sample, stranded TruSeq cDNA libraries were prepared using poly(A) enrichment. Libraries were sequenced on an Illumina NovaSeq 5000 platform to generate 150 bp paired-end (PE) reads.

The raw reads were processed using Fastp (v.0.23.4) (Chen et al., 2018) with default trimming options and base correction enabled to correct for low-quality bases. Processed paired-end reads were then quantified using Salmon (v1.10.3) (Patro et al., 2017). A Salmon index was built from the reference *C. reinhardtii* v.6.1 transcriptome downloaded from Phytozome. Quantification was performed with automatic library type detection (l A), validation of mappings (validateMappings), and corrections for both sequence-specific (seqBias) and GC (gcBias) biases. One hundred bootstraps (numBootstraps 100) were performed for each sample.

All further analysis was carried out in Rstudio (v.2024.04.1, Posit team, 2023). The R tximport package (v.1.32.0) (Soneson et al., 2015) was used to import the resulting gene-level expression estimates. Genes with read counts fewer than 10 reads in at least six samples were marked as lowly expressed and removed from further analysis.

Differential expression analysis was carried out using DESeq2 package (v.1.44.0) (Love et al., 2014). A gene was considered statistically significant if the |log_2_ FC| was ≥ 1 and the adjusted *P*-value < 0.01. Cultures grown at 18 °C served as the reference condition for all pairwise comparisons. Genes with positive log₂ fold changes were designated as upregulated, and those with negative values were designated as downregulated relative to the 18 °C condition. Gene ontology enrichment was carried out using topGO (v.2.56.0). To compare different metabolic pathways, Chlamydomonas genes were assigned to specific pathways using the *C. reinhardtii* vs 6.1 KEGG annotations and the KEGG Reconstruct tool. Transcript abundances of genes belonging to identified pathways were then visualized using heatmaps.

### Isolation of flagella and ciliary proteins

Cells were grown under the above-mentioned conditions for four days at 18 °C and 28 °C, respectively. Cells were harvested and washed according to Bösger et al. (2014). The temperature of rotors and washing buffers was adjusted to 18 °C and 28 °C, respectively. After deflagellation of the cells with dibucain and purification of the flagella using sucrose step-gradient centrifugation, the ciliary proteins were prepared by adding NP-40 to the isolated flagella to achieve a final concentration of 1%, as described in Bösger et al. (2014). NP-40 is used as mild detergent to solubilize membrane proteins. Four independent biological replicates were prepared for each temperature condition, from which three were used for further analysis.

### Proteome analysis of the NP-40 treated ciliary proteins

For the analysis of the ciliary proteins, 0.5 mg proteins of each experiment were reduced and carbamidomethylated according to Zhang et al., 2005. Samples were then subjected to the single-pot, solid phase-enhanced sample preparation (SP3) according to Hughes et al., (2019). Briefly, the proteins were bound to magnetic SP3 beads, washed three times to remove contaminants, and digested overnight at 37 °C directly on the beads by adding 20 µg of a Trypsin and LysC protease mix (MS grade, Promega V507A) dissolved in 40 µL of the provided buffer per 0.5 mg sample in Eppendorf Protein LoBind tubes that were shaken at 1,000 rpm in a Thermomixer. Prior to the mass spectrometric analyses, the resulting peptides were further purified by reversed phase chromatography using ZipTip microtips supplemented with additional Poros R2 10 μm reversed phase material (Bösger et al., 2009), dried using a Speed-Vac and stored at – 80 °C before mass spectrometry analysis. Immediately before LC-ESI-MS/MS, peptides were reconstituted in 25 µL of 5% MeCN/0.1% TFA by vortexing thrice and a 5 min cycle of ultrasonic bath sonication. The samples were then centrifuged for 5 min at 13,000 rpm and the supernatant was transferred to MS vials for LC-MS/MS measurements.

### Preparation of samples for the secretory proteome

Secretome analysis was performed using cultures grown at 18 °C and 28 °C. Cells were cultivated for four days in TAP medium, as described above. Three independent biological replicates were prepared for each temperature condition. On day 4, cultures were harvested to collect cell-free supernatants. For each sample, the culture was centrifuged at 1,000 × *g* for 10 min to remove intact cells and large particulates. The resulting supernatant was then filtered through a 0.22 µm pore-size membrane filter (Carl Roth, Germany) to ensure complete removal of any residual cells or debris.

10 mL of the supernatant from each sample were first concentrated to 250 µL using a 3 kDa MWCO Amicon centrifugal filter. Each concentrate was transferred into an Eppendorf Protein LoBind tube and adjusted to 15 µg total protein, as determined by BCA assay. The samples were finally brought to 250 µL with 1 M triethyl ammonium bicarbonate (TEAB) buffer, yielding a final concentration of 50 mM TEAB (pH ∼8.5), and incubated at 60 °C for 10 min with shaking at 1,000 rpm.

The subsequent steps were performed at 22 °C with shaking at 1,500 rpm. Proteins were sequentially reduced by adding 2.5 µL TCEP to a final concentration of 5 mM and incubated for 10 min, then alkylated with 5 µL iodoacetamide to a final concentration of 10 mM for 30 min and finally quenched by adding 5 µL DTT to a final concentration of 10 mM for 15 min. Next, a 1:1 mixture of hydrophobic and hydrophilic Sera-Mag carboxylate coated magnetic beads (50 mg mL^-1^; 2 mg total; particles E3/E7; Cytiva) were prewashed three times with 100 µL water. After washing, 2.5 µL of the bead suspension was added to each sample, followed by ethanol to a final concentration of 60%. Samples were incubated for 5 min to bind proteins. Beads were bound using a magnetic rack. The supernatant was removed, and the beads were washed three times with 400 µL of 80% ethanol and once with 400 µL of MeCN. After an air-drying step, the beads were resuspended in 50 µL of 50 mM TEAB (pH ∼8.5).

Proteins on the beads were digested by adding 0.5 µL of recombinant LysC (0.2 µg µL^-1^, MS grade, Promega V1671) and incubation at 37 °C for 1 h with shaking at 1,200 rpm. Subsequently, 0.4 µL of trypsin (0.5 µg µL^-1^, MS grade, Promega VA9000) was added, and digestion continued overnight at 37 °C with shaking at 1,200 rpm. After digestion, the supernatant was collected into a new Protein LoBind tube. The beads were washed twice with 20 µL of 1% TFA, whereby each wash was incubated for 5 min at 22 °C while shaking at 1,000 rpm. The combined supernatant and wash fractions were acidified with 1 µL of 80% TFA (to pH < 3) and desalted using a 100 µL Pierce C18 Tips (Thermo Fisher 87784). Tips were equilibrated with 100 µL of 50% MeCN/0.1% TFA twice, followed by equilibration with 100 µL of 0.1% TFA twice. Samples were loaded by slowly aspirating and dispensing the solution ten times. Tips were washed twice with 100 µL of 0.1% TFA, and bound peptides were eluted three times with 75 µL of 80% MeCN/0.1% TFA. Eluates were combined, dried in a vacuum concentrator at 35 °C, and stored at –80 °C until analysis. Immediately before LC-ESI-MS/MS, peptides were reconstituted in 25 µL of 5% MeCN/0.1% TFA by vortexing thrice and a 5 min cycle of ultrasonic bath sonication. The samples were then centrifuged for 5 min at 13,000 rpm and the supernatant was transferred to MS vials for LC-MS/MS measurements.

### LC-ESI-MS/MS measurements

LC-MS/MS analysis was performed with an Ultimate3000 Nano-HPLC system (Thermo Fisher Scientific) coupled to an Orbitrap Fusion Tribrid mass spectrometer (Thermo Fisher Scientific). Samples were analyzed using an Acclaim PepMap 100 C18 trap column (0.075 × 200 mm; Thermo Fisher Scientific) and an Aurora Ultimate XT C18 UHPLC analytical emitter column (0.075 × 250 mm; Ionopticks, Collingwood, Australia) in an EASY Spray source setting with HeatSync Column Heater (Ionopticks, Collingwood, Australia). Each sample was loaded onto the trap column, washed with 0.1% TFA in water, and then transferred to the analytical column. Peptides were separated in a 152-minute run (buffer A: water with 0.1% FA, buffer B: MeCN with 0.1% FA, flowrate: 400 nL/min) with the following gradient: from 5% B to 22% B in 105 min, from 22% B to 32% B in 10 min, from 32% B to 90% B in 10 min, hold at 90% B for 10 min, then to 5% B over 0.1 min, and hold at 5% B for 10 min. The Orbitrap Fusion Tribrid mass spectrometer was operated in a data-dependent acquisition (DDA) mode with a cycle time of 3 s. Full scan acquisition was performed in the Orbitrap at a resolution of 120,000 in the scan range of 300 – 1,500 m/z (AGC target of 50%, maximal injection time of 50 ms). Monoisotopic precursor selection as well as dynamic exclusion (exclusion duration: 60 s) was enabled. Precursors with charge states of 2 - 7 and intensities greater than 5e3 were selected for fragmentation. Isolation was performed in the quadrupole using a window of 1.6 m/z. Fragments were generated using higher-energy collisional dissociation (HCD, normalized collision energy: 30%) and detected in the Ion Trap (scan rate: rapid, mass range: normal, maximal injection time: 35 ms, AGC target: standard).

### Proteomics data analysis along with label-free quantification

All raw LC–MS/MS data were processed with FragPipe version (v22.0), incorporating MSFragger (v4.1) for peptide identification. We applied the built in LFQ MBR workflow to search against the publicly available Chlamydomonas v6.1 reference proteome database supplemented with common contaminants and decoy sequences (CreinhardtiiCC_4532_707_v6.1.protein_primaryTranscriptOnly.fa). MSFragger performed a closed search using initial precursor and fragment mass tolerances of 0.5 Da, followed by automatic mass recalibration and selection of narrower tolerances. The enzyme was set to strict trypsin, allowing up to two missed cleavages, with peptide lengths constrained to 7–50 residues and masses between 500 and 5,000 Da. Methionine oxidation (+15.9949 Da) and protein N-terminal acetylation (+42.0106 Da) were set as variable modifications, while cysteine carbamidomethylation (+57.0215 Da) was fixed. Subsequently, MSBooster, Percolator, and Philosopher were employed to generate deep learning augmented scores, perform PSM rescoring, and estimate false discovery rates. We implemented a dynamic filtering threshold based on 2.5% of the global maximum peptide count per protein. Thus, the inclusion cutoffs were set to at least eight unique peptides per protein for the ciliary proteome (maximum peptide count per protein: 304) and to at least three unique peptides per protein for the secretome (maximum peptide count per protein: 110). Label-free quantification was then carried out with IonQuant (v1.10.27) under default settings, using a mass tolerance of 10 ppm and a retention time tolerance of 0.4 min. Both, “match between runs” and “MaxLFQ” algorithms were enabled. The “min scans” parameter was set to 3, “min isotopes” was set to 2, and “MaxLFQ min ions” was set to 2. Proteins were considered differentially regulated when they exhibited a |log_2_ fold change| ≥ 0.5 and an adjusted *P*-value < 0.05.

### Databases used for the determination of ciliary proteins

For ciliary protein identification, the Chlamydomonas ciliary protein database established by Pazour et al., 2005 (https://chlamyfp.org/ChlamyFPv2/cr_read_sql.php) was used as a default. The resulting candidates are labeled in dark blue in Supplementary Tables 9, 10 and 11. Moreover, the cilia proteinatlas (https://www.proteinatlas.org/search/cilia) database referenced by Hansen et al., 2025 that combines data from ciliary proteins of mouse, humans, bovine and Xenopus was used. For this purpose, the provided XML file was downloaded and manually searched for the candidates. In addition, the primary cilium proteome database (https://esbl.nhlbi.nih.gov/Databases/CiliumProteome/) compiled from Ishikawa et al., 2012, Mick et al., 2015 and May et al., 2021 was used. Candidates deriving from these databases or being directly found by publications are labeled in light blue in Supplementary Tables 9, 10 and 11.

### Predictions of extracellular proteins

DeepLoc 2.1 (Odum et al., 2024) was used to predict the subcellular localization of identified secreted proteins, with the organism type set to “Plant” to optimize predictions for the Chlorophyte alga Chlamydomonas. DeepLoc 2.1 predicts localization up to ten subcellular compartments: nucleus, cytoplasm, extracellular/secreted, mitochondrion, cell membrane, endoplasmic reticulum, chloroplast, Golgi apparatus, lysosome/vacuole, and peroxisome. Proteins were classified as secreted if they received an “extracellular” prediction with a confidence score > 0.5.

### Evans blue staining of cells

200 µL of algal cultures grown at 18 °C and 28 °C for four days were mixed during day-phase (LD4) with 200 µL of 0.2% (w/v) Evans Blue in TAP, resulting in a final dye concentration of 0.1% (w/v), as described by Hotter et al. (2021). The experiments were performed using two independent datasets, each with three biological replicates. Each biological replicate was divided into three technical replicates. After 5 min incubation at room temperature, the cells were washed twice with TAP by centrifugation at 11,000 × *g* and resuspended in 180 µL TAP medium. 20 µL of Lugol’s solution was added to immobilize the algal cells, and the percentage of stained cells was determined using brightfield microscopy (Axiophot, Carl Zeiss, Germany). For each technical replicate, 50 cells were examined.

### Determination of acetate levels

1 mL of algal cultures grown at 18 °C and 28 °C, respectively, and harvested during day-phase (LD4) by centrifugation at 11,000 × *g* for 5 min at 4 °C. For each sample, 700 μl of the top clear supernatant was collected into a new tube. The supernatant was stored in a freezer at –80 °C until analysis. The acetate content in the supernatant was quantified using an acetic acid assay kit (Megazyme, Wicklow, Ireland) following the manufacturer’s instructions. The experiments were performed using two independent datasets, each along with three biological replicates, each of them containing three technical replicates.

### Determination of CO_2_ levels in water

The CO₂ concentration in water was measured using the CA–carbon dioxide test kit (Hach Lange GmbH, Berlin, Germany). For this purpose, 23 mL of water was dispensed into Nunc flasks and incubated at 18 °C, 23 °C, 28 °C, and 33 °C, respectively, for 24 hours prior to measurement. CO₂ levels were determined in triplicate using the kit according to the manufacturer’s instructions.

### CO_2_ transfer rate (CTR) measurements

An ISF1-Z compressor shaking incubator (Adolf Kühner AG, Birsfelden, Switzerland) was taken for the experiment. CTR measurements were carried out using algal precultures grown either at 18 °C or 28 °C. Three biological replicates with independent precultures were performed for each temperature. Cultures were adjusted to an initial density of 1 × 10^5^ cells mL⁻¹ in TAP medium and put into 250 mL Erlenmeyer flasks suited for CO_2_ measurements. Cells were incubated at 18 °C and 28 °C, respectively, at 100 rpm. The orbital shakers were equipped with the Kuhner Transfer-Rate Online Measurement (TOM) system for off-gas analysis and online carbon dioxide transfer rate (CTR) monitoring, and a PHOTOBioSim light module with a control unit (Adolf Kühner AG, Birsfelden, Switzerland). Data were taken every 60 min over four days. One off-gas measuring cycle of 60 min included 20 min measuring time. The needed aeration rate with compressed air was set to 10 mL min^-1^. Cells were exposed, as for all other growth studies, to a light-dark cycle of 12 h light and 12 h darkness. For the light period, white light with a full spectrum was used, as before, and the light intensity was set to approximately 50 µmol m^-2^ s^-1^.

### Crude extracts and immunoblots

Crude extracts of soluble proteins (Zhao et al., 2004) were used for CRY-DASH1, CRY-DASH2 and aCRY detection and total protein extracts for CHR1 detection (Vuong et al., 2025). Cells were harvested at LD4. For tubulin and CK1 detection, NP-40 treated ciliary proteins were used. Isolation of flagella and ciliary proteins was performed as described above. Antibodies against CRY-DASH1 (Rredhi et al., 2021), aCRY (Beel et al., 2012), CHR1 (Vuong et al., 2025) and CK1 (Schulze et al., 2013) were used. Antibodies against acetylated α-tubulin were obtained from Sigma-Aldrich (T7451), and antibodies against α-tubulin from Agrisera (AS10 680). Antibodies against CRY-DASH2 were produced by the Pineda Antikörper-Service (Berlin) using full-length CRY-DASH2 protein with an additional amino acid (glutamic acid) after the first methionine and a 6x His-tag at its C-terminus. For this purpose, the *CRY-DASH2* coding sequence was codon-adapted for *Escherichia coli* and synthesized by GeneArt within plasmid 13AA7FKP-CDS294-pMK-RQ (Supplementary Figure 14). The according gene was cloned in vector pET28a(+) with restriction sites *Nco*I and *Hind*III. His-tagged CRY-DASH2 was overexpressed in *E. coli* and affinity purified as described for CRY-DASH1 (Rredhi et al., 2021). Immunoblots were performed as described previously (Vuong et al., 2025). Briefly, proteins were separated by SDS-PAGE and transferred to polyvinylidene difluoride (PVDF) membranes or nitrocellulose membranes (for anti-CRY-DASH2 antibodies) that were stained with Coomassie Brilliant Blue R 250 or Ponceau S (for anti-CRY-DASH2 antibodies). The following dilutions of primary antibodies were taken: 1:5,000 for anti-CRY-DASH1 and anti-aCRY, 1:1,000 for anti-CRY-DASH2, 1:2,000 for anti-CHR1, 1:5,000 for CK1, 1:10,000 for anti-acetylated α-tubulin (T7451, Sigma-Aldrich) and 1:20,000 for anti-α-tubulin (AS10 680, Agrisera). The horseradish peroxidase-conjugated anti-rabbit IgGs (Sigma-Aldrich) were used as the secondary antibody for the photoreceptors, α-tubulin and CK1, while horseradish peroxidase-conjugated anti-mouse IgGs (Sigma-Aldrich) were implemented for acetylated α-tubulin. Dilutions of the secondary antibody were 1:5,000 (CHR1, CK1, CRY-DASH2, and aCRY), 1:6,666 (CRY-DASH1), 1:10,000 (acetylated α-tubulin), and 1:20,000 (α-tubulin).

### Determination of flagellar length

Flagellar length was quantified on days 1, 3, 4, and 7 of cell growth. 2 mL aliquots from the main Chlamydomonas cultures were harvested and centrifuged at 500 × *g* for 10 min. Ciliary immunolocalization studies were done according to Zou et al. (2017). Cells were incubated with a primary antibody against acetylated α-tubulin (Sigma, clone 6-11B-1, mouse monoclonal) at a dilution of 1:5,000, followed by a goat anti-mouse secondary antibody conjugated to AlexaFluor 555 (Life Technologies, A32727) at a 1:2,000 dilution. Cilia were visualized using Zeiss Axiophot and Leica Stellaris 8 Fluorescent Microscopes, respectively. Flagella length was measured with ImageJ (version 1.53t, Schneider et al., 2012). Statistical analyses and graphing were conducted in RStudio (v.2024.04.1, Posit team, 2023).

### Motility analysis

Motility analysis of algal cells was performed using day 4 cultures grown in TAP medium at the different temperatures, whose cell density had been standardized to 5 × 10^5^ cells mL^-1^. A 120 µL aliquot was transferred onto a sterile glass microscope slide fitted with a Gene Frame adhesive chamber (Thermo Fisher Scientific, catalog no. AB0578). Prior to placing the Gene Frame adhesive chamber, slides were cleaned with ethanol and sterilized under UV light for 15 min. After the sample was added, the chamber was sealed with a coverslip that had also been pre-cleaned with ethanol and UV-sterilized. Live-cell imaging was conducted using an inverted Nikon Ti2 widefield microscope under a 4x/0.13 NA Plan Fluor PhL DL (Nikon, Japan) objective. Images were captured with a Hamatsu C 14440 – 20UP camera. Temperature control during imaging was achieved using a cooling/heating Incubation Insert P-Set 2000 (PeCon, Germany) coupled to a DYNEO DD-310F refrigerated and heating circulator (JULABO GmbH, Germany). The imaging temperature was maintained at either 18 °C or 28 °C to match the cultivation conditions of each sample. Time-lapse recordings were acquired at intervals of 200 ms per frame. Image sequences were subsequently imported into ImageJ to perform brightness and contrast correction, background subtraction and image thresholding. This results in a series of binary images in which Chlamydomonas cells can be clearly distinguished from the background. Individual cell trajectories were tracked using the TrackMate plugin, which quantified key motility parameters including turning frequency and swimming speed. The extracted tracking data were then exported for statistical analysis and visualization. Trajectory plots were visualized in RStudio.

To assess thermal acclimation, cells grown for four days at 18 °C were transferred to a water bath pre-set to 28 °C and incubated for 15 min. After this incubation, slides were prepared as described above. Imaging was performed at a maintained temperature of 28 °C.

### Gametogenesis and mating ability assay

(Jiang and Stern, 2009) Chlamydomonas vegetative cells of strains CC-125 (mt^+^) and CC-124 (mt^−^) were first transferred from liquid TAP cultures onto solid TAP agar plates and incubated for 4 days at either 18 °C or 28 °C under a 12 h:12 h LD cycle. Following this, cells were transferred onto N10 agar plates and incubated in the dark at their respective temperatures for three additional days to induce nitrogen starvation and promote pregamete formation. Following this, cells were harvested from the N10 plates and resuspended in 10 mL of sterile distilled water that had been acclimatized to the corresponding temperature. Cell suspensions were transferred to sterile NUNC flasks and placed under continuous intense illumination at 70 µmol m^-2^ s^-1^ for 2–3 h without agitation to induce gamete formation. Following induction, mt^+^ and mt^-^ gametes were adjusted to a final concentration of 2 × 10^6^ cells mL^-1^ using a hemocytometer. 1 mL from each mating type were combined in fresh NUNC flasks and kept under continuous light without shaking to allow mating. Mating efficiency was evaluated 1 h after mixing the gametes. Samples from each flask were collected and counted using a light microscope. Large, fused cells were counted as single units. Mating efficiency was calculated using the formula:

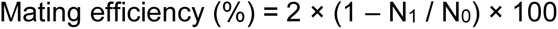

In this equation, N_0_ represents the number of cells per mL immediately after mixing, and N_1_ is the number of cells per mL measured 1 h after incubation (Chiang et al., 1970).

### Accession numbers

Transcriptome data from this article can be found under BioProject PRJNA1306621. Motility videos are deposited to Zenodo under DOI: 10.5281/zenodo.16885208. Proteome data was uploaded to PRIDE PXD067459 and contain the full list of total peptides from Chlamydomonas as well as other contaminants (e.g. trypsin, keratin). Custom scripts for the transcriptome, proteome and motility data are provided in GitHub under the following link: https://github.com/HitMonk/Ambient_temperature_multiomics

## Supplementary Data

**Data Set 1:**

**Supplementary Figure 1.** Boxplot analysis of cell lengths and estimated cell areas measured at LD6

**Supplementary Figure 2**. Cell length distributions at different ambient temperatures and cultivations days measured at LD6

**Supplementary Figure 3**. Hierarchical clustering of all differentially abundant transcripts from cells grown at 18 °C versus 28 °C

**Supplementary Figure 4.** Temperature-dependent regulation of transcripts encoding TRP channels

**Supplementary Figure 5.** Growth curves of Chlamydomonas mono- and cocultures with *P. protegens* at 23 °C (A) and 33 °C (B)

**Supplementary Figure 6.** Expression profiles of transcripts encoding RNA-binding proteins

**Supplementary Figure 7.** Heatmap of transcript abundances at different ambient temperatures encoding proteins of photosynthesis and porphyrin metabolism (chlorophyll biosynthesis)

**Supplementary Figure 8.** Effects of temperature on CO_2_ levels

**Supplementary Figure 9.** Flagellar length distributions at different ambient temperatures on day 4 at LD6

**Supplementary Figure 10.** Quantification of flagellar length in cells grown at different temperatures on days 1, 3, and 7

**Supplementary Figure 11.** Cell movements with mean directional change and median swimming speed (independent replicates)

**Supplementary Figure 12.** Mating efficiency at different temperatures (second independent dataset)

**Supplementary Figure 13.** Evans Blue staining of cells grown at 18 °C or 28 °C for four days

**Supplementary Figure 14.** Codon adapted *CRY-DASH2*

**Data Set 2:**

**Supplementary Table 1.** Values for growth kinetics of Chlamydomonas at different ambient temperatures compared to 23 °C

**Supplementary Table 2A, B.** (A) Values for cell lengths at different ambient temperatures compared to 18 °C; (B) Values for estimated cell areas at different ambient temperatures compared to 18 °C

**Supplementary Table 3.** Transcript abundances altered in cells grown at different ambient temperatures

**Supplementary Table 4.** Full list of gene ontology (GO) terms of differentially expressed transcripts from cells cultivated at 28 °C and 18 °C

**Supplementary Table 5.** List of selected candidates from transcriptome and proteome studies along with the genome model numbers from Figures 2 to 6 and Supplementary Figure 4, 6, and 7

**Supplementary Table 6.** OD values of *P. protegens* bacteria grown at different ambient temperatures

**Supplementary Table 7.** List of selected candidates along with the genome model numbers from metabolic pathways that are not altered in a coordinated way upon ambient temperature changes

**Supplementary Table 8.** List of CO_2_ transfer rates (data points measured every 60 min over four days) for three biological replicates from cells grown at 18 °C or 28 °C **Supplementary Table 9.** List of identified proteins from NP-40 treated cilia isolated from cells grown at 18 °C or 28 °C after application of a 2.5% peptide count filter (at least eight unique peptides)

**Supplementary Table 10.** List of identified ciliary proteins from cells grown at 18 °C or 28 °C after application of a 2.5% peptide count filter (at least eight unique peptides)

**Supplementary Table 11.** List of temperature-dependent significantly up- and downregulated ciliary proteins (at least eight unique peptides)

**Supplementary Table 12.** List of ciliary lengths of cells grown at different ambient temperatures compared to 18 °C

**Supplementary Table 13.** List of identified secretory proteins from cells grown at 18 °C or 28 °C after application of a 2.5% peptide count filter (at least three unique peptides)

**Supplementary Table 14.** List of identified secretory proteins determined by DeepLoc (at least three unique peptides)

**Supplementary Table 15.** List of temperature-dependent significantly up- and downregulated secretory proteins (at least three unique peptides)

**Video 1.** Motility patterns of Chlamydomonas cells grown at 18 °C

**Video 2.** Motility patterns of Chlamydomonas cells grown at 28 °C

## ACKNOWLEDGMENTS

We thank Aurelie Jost from the Microverse Imaging Center for help with the Leica Stellaris 8 Fluorescence Microscope (DFG project ID 460889961), Lysett Wagner for support with cloning and Melvin Schubert for help with overexpression of CRY-DASH2. PS, MAR, RJA, CCP, ML and MM received funding from the Deutsche Forschungsgemeinschaft (DFG, German Research Foundation) within the Germany’s Excellence Strategy – EXC 2051 – Project ID 390713860. JC was funded by DFG under CRC 1127, Project ID 239748522. CL was supported by a PhD grant of the China Scholarship Council (CSC). ML is grateful for support by the research profile line LIFE of FSU Jena and the Deutsche Forschungsgemeinschaft (DFG, German Research Foundation) via the Emmy-Noether-Program (Project ID 528114058) and CRC 1127, Project ID 239748522. We thank the Microverse Imaging Center also financed by EXC 2051 for providing microscope facility support for data acquisition.

## AUTHOR CONTRIBUTIONS

PS, TV, MAR, RJA, ML and MM designed the project. PS, TV, CL, VW, DM, CCP, WL, JC, AZ and SW performed experiments, and all authors contributed to analysis and interpretation of results. PS, TV and MM wrote the paper with input from all coauthors.

## Notes

### Competing Interest Statement

The authors have declared no competing interest.

### Summary of Updates

Updated the manuscript title and added new authors to reflect their contributions to newly performed experiments. Extended methods. Added boxplots showing cell and flagella lengths at days 1 and 7, as well as cell areas at days 1 and 4. Included supplementary plots on cell length and flagella length distributions Reanalyzed the transcriptome data using Salmon instead of the Kallisto program. Added growth curve data for Pseudomonas protegens at various temperatures. Included immunoblots for additional photoreceptors across three different temperatures. Performed a transcriptome analysis of metabolic pathways using KEGG mapping and pathway codes. Provided data on CO2 transfer rates, as well as CO2 and acetate levels, at different temperatures. Added immunoblots against alpha-tubulin. Included data on motility changes in response to short-term temperature shifts. Added data addressing potential cell lysis at different temperatures. Applied a peptide count threshold to the ciliary and secretory proteomes. Conducted a manual inspection of ciliary proteins against additional ciliary databases and literature.

## REFERENCES

Aiyar, P, Schaeme, D, García-Altares, M, Carrasco Flores, D, Dathe, H, Hertweck, C, Sasso, S and Mittag, M (2017) Antagonistic bacteria disrupt calcium homeostasis and immobilize algal cells. Nature Communications, 8: 1756

Alexandersson, E, Ali, A, Resjö, S and Andreasson, E (2013) Plant secretome proteomics. Frontiers in Plant Science, 4: 9

Barolo, L, Commault, AS, Abbriano, RM, Padula, MP, Kim, M, Kuzhiumparambil, U, Ralph, PJ and Pernice, M (2022) Unassembled cell wall proteins form aggregates in the extracellular space of *Chlamydomonas reinhardtii* strain UVM4. Applied Microbiology and Biotechnology, 106: 4145–4156

Barrett, MR and Koch, AR (1982) Effects of ammonium sulfate and urea on the growth of Chlorophycean algae from rice fields. Journal of Environmental Quality, 11: 187–191

Beel, B, Prager, K, Spexard, M, Sasso, S, Weiss, D, Müller, N, Heinnickel, M, Dewez, D, Ikoma, D, Grossman, AR, et al. (2012) A flavin binding cryptochrome photoreceptor responds to both blue and red light in Chlamydomonas reinhardtii. Plant Cell 24: 2992–3008

Bösger, J, Wagner, V, Weisheit, W and Mittag, M (2009) Analysis of flagellar phosphoproteins from *Chlamydomonas reinhardtii*. Eukaryotic Cell, 8: 922–932

Bösger, J, Wagner, V, Weisheit, W and Mittag, M (2014) Comparative phosphoproteomics to identify targets of the clock-relevant casein kinase 1 in *C. reinhardtii* flagella. Methods in Molecular Biology, 1158: 187–202

Bunbury, F, Deery, E, Sayer, AP, Bhardwaj, V, Harrison, EL, Warren, MJ and Smith, AG (2022) Exploring the onset of B_12_-based mutualisms using a recently evolved *Chlamydomonas* auxotroph and B_12_-producing bacteria. Environmental Microbiology, 24: 3134–3147

Burgunter-Delamare, B, Shetty, P, Vuong, T and Mittag, M (2024) Exchange or eliminate: The secrets of algal-bacterial relationships. Plants, 13: 829

Carrasco Flores, D, Hotter V, Vuong T, Hou Y, Bando Y, Scherlach K, Burgunter-Delamare B, Hermenau R, Komor AJ, Aiyar P et al. (2024) A mutualistic bacterium rescues a green alga from an antagonist. Proceedings of the National Academy of Sciences USA of the United States of America, 121: e2401632121

Catalan, RE, Fragkopoulos, AA, Girot, A, Lorenz, M and Bäumchen, O (2025) Preparation, maintenance and propagation of synchronous cultures of photoactive *Chlamydomonas* cells. Nature Protocols, 20: 2125–2150

Chen, S, Zhou, Y, Chen, Y and Gu, J (2018) fastp: an ultra-fast all-in-one FASTQ preprocessor. Bioinformatics, 34: i884–i890

Chiang, KS, Kates, JR, Jones, RF and Sueoka, N (1970) On the formation of a homogeneous zygotic population in *Chlamydomonas reinhardtii*. Developmental Biology, 22: 655–669

Chin, KJ, Lukow, T and Conrad, R (1999) Effect of temperature on structure and function of the methanogenic archaeal community in an anoxic rice field soil. Applied and Environmental Microbiology, 65: 2341–2349

Christie, JM and Zurbriggen, MD (2021) Optogenetics in plants. New Phytologist, 229: 3108–3115

Craig, RJ, Gallaher, SD., Shu, S, Salomé, PA, Jenkins, JW, Blaby-Haas, CE, Purvine, SO, O’Donnell, S, Barry, K, Grimwood, J, et al. (2023) The Chlamydomonas Genome Project, version 6: Reference assemblies for mating-type plus and minus strains reveal extensive structural mutation in the laboratory. The Plant Cell, 35: 644–672

Crockett, C (2021) How a microorganism corkscrews while it breaststrokes. Physics, 14: s21

Crozet, P, Navarro, FJ, Willmund, F, Mehrshahi, P, Bakowski, K, Lauersen, KJ, Pérez-Pérez, ME, Auroy, P, Gorchs Rovira, A, Sauret-Gueto, S and Niemeyer, J, (2018) Birth of a photosynthetic chassis: a MoClo toolkit enabling synthetic biology in the microalga *Chlamydomonas reinhardtii*. ACS Synthetic Biology, 7: 2074–2086

Deisseroth, K and Hegemann, P (2017) The form and function of channelrhodopsin. Science, 357: eaan5544

Dupuis, S and Merchant, SS (2023) *Chlamydomonas reinhardtii*: A model for photosynthesis and so much more. Nature Methods, 20: 1441–1442

Ermilova, E (2020) Cold stress response: an overview in *Chlamydomonas*. Frontiers in Plant Science, 11: 569437

Folcik, AM, Cutshaw, K, Haire, T, Goode, J, Shah, P, Zaidi, F, Richardson, B and Palmer, A (2020) Quorum sensing behavior in the model unicellular eukaryote Chlamydomonas reinhardtii. iScience, 23: e101835

Ghasemi Y, Rasoul-Amini S, Morowvat MH, Raee MJ, Ghoshoon MB, Nouri F, Negintaji N, Parvizi R, Mosavi-Azam SB. (2008) Characterization of hydrocortisone biometabolites and 18S rRNA gene in *Chlamydomonas reinhardtii* cultures. Molecules, 13: 2416–2425

Greiner, A, Kelterborn, S, Evers, H, Kreimer, G, Sizova, I and Hegemann, P (2017) Targeting of photoreceptor genes in *Chlamydomonas reinhardtii* via zinc-finger nucleases and CRISPR/Cas9. The Plant Cell, 29: 2498–2518

Hallmann, A (2006) The pherophorins: common, versatile building blocks in the evolution of extracellular matrix architecture in Volvocales. Plant Journal, 45: 292–307

Hansen, JN, Sun, H, Kahnert, K, Westenius, E, Johannesson, A, Villegas, C, Le, T, Tzavlaki, K, Winsnes, C, Pohjanen, E and Mäkiniemi, A et al. (2025) Intrinsic heterogeneity of primary cilia revealed through spatial proteomics. Cell, 188: 6804–6824

Harris, EH (1989) The Chlamydomonas sourcebook: a comprehensive guide to biology and laboratory use. Academic Press, Cambridge

Helliwell, KE, Collins, S, Kazamia, E, Purton, S, Wheeler, GL, and Smith, AG (2015) Fundamental shift in vitamin B_12_ eco-physiology of a model alga demonstrated by experimental evolution. The ISME Journal, 9: 1446–1455

Hey, S and Linder, S (2024) Matrix metalloproteinases at a glance. Journal of Cell Science, 137: jcs261898

Hori, T, Noll, M, Igarashi, Y, Friedrich, MW, and Conrad, R (2007) Identification of acetate-assimilating microorganisms under methanogenic conditions in anoxic rice field soil by comparative stable isotope probing of RNA. Applied and Environmental Microbiology, 73: 101–109

Hou, Y, Bando, Y, Carrasco Flores, D, Hotter, V, Das, R, Schiweck, B, Melzer, T, Arndt, HD and Mittag, M (2023) A cyclic lipopeptide produced by an antagonistic bacterium relies on its tail and transient receptor potential-type Ca^2+^ channels to immobilize a green alga. New Phytologist, 237: 1620–1635

Hotter, V, Zopf D, Kim HJ, Silge A, Schmitt M, Aiyar P, Fleck J, Matthäus C, Hniopek J, Yan Q, et al. (2021) A polyyne toxin produced by an antagonistic bacterium blinds and lyses a Chlamydomonad alga. Proceedings of the National Academy of Sciences of the United States of America, 118: e2107695118

Hughes, CS, Moggridge, S, Müller, T, Sorensen, PH, Morin, GB and Krijgsveld, J (2019) Single-pot, solid-phase-enhanced sample preparation for proteomics experiments. Nature Protocols, 14: 68–85

Ishikawa, H, Thompson, J, Yates III, JR, and Marshall, WF (2012) Proteomic analysis of mammalian primary cilia. Current Biology, 22: 414–419

Jain, N, Lim, LW, Tan, WT, George, B, Makeyev, E and Thanabalu, T (2014) Conditional N-WASP knockout in mouse brain implicates actin cytoskeleton regulation in hydrocephalus pathology. Experimental Neurology, 254: 29–40

Jeffery, CJ (2018) Protein moonlighting: what is it, and why is it important?. Philosophical Transactions of the Royal Society B: Biological Sciences, 373: 1738

Jiang, X and Stern, D (2009) Mating and tetrad separation of *Chlamydomonas reinhardtii* for genetic analysis. Journal of Visualized Experiments, 30: 1274

Katoh, Y, Terada, M, Nishijima, Y, Takei, R, Nozaki, S, Hamada, H and Nakayama, K (2016) Overall architecture of the intraflagellar transport (IFT)-B complex containing Cluap1/IFT38 as an essential component of the IFT-B peripheral subcomplex. Journal of Biological Chemistry, 291: 10962–10975

Kinoshita, T, Fukuzawa, H, Shimada, T, Saito, T and Matsuda, Y (1992) Primary structure and expression of a gamete lytic enzyme in *Chlamydomonas reinhardtii*: similarity of functional domains to matrix metalloproteases. Proceedings of the National Academy of Sciences of the United States of America, 89: 4693–4697

Lacey, SE, Graziadei, A and Pigino, G (2024) Extensive structural rearrangement of intraflagellar transport trains underpins bidirectional cargo transport. Cell, 187: 4621–4636

Lechtreck, KF, Johnson, EC, Sakai, T, Cochran, D, Ballif, BA, Rush, J, Pazour, GJ, Ikebe, M and Witman, GB (2009) The *Chlamydomonas reinhardtii* BBSome is an IFT cargo required for export of specific signaling proteins from flagella. Journal of Cell Biology, 187: 1117–1132

Légeret, B, Schulz-Raffelt, M, Nguyen, HM, Auroy, P, Beisson, F, Peltier, G, Blanc, G, Li-Beisson, Y (2016) Lipidomic and transcriptomic analyses of *Chlamydomonas reinhardtii* under heat stress unveil a direct route for the conversion of membrane lipids into storage lipids. Plant, Cell & Environment, 39: 834–847

L’Hernault, SW and Rosenbaum, JL (1985) Chlamydomonas α-tubulin is posttranslationally modified by acetylation on the ε-amino group of a lysine. Biochemistry, 24: 473–478

Li, W, Flores, DC, Füßel, J, Euteneuer, J, Dathe, H, Zou, Y, Weisheit, W, Wagner, V, Petersen, J and Mittag, M (2018) A musashi splice variant and its interaction partners influence temperature acclimation in *Chlamydomonas*. Plant Physiology, 178: 1489–1506

Li, X, Liu, Q and Pan, J (2025) Chlamydomonas as a model system for the study of cilia and eukaryotic flagella. Seminars in Cell & Developmental Biology, 175: 103658

Li, X, Patena, W, Fauser, F, Jinkerson, RE, Saroussi, S, Meyer, MT, Ivanova, N, Robertson, JM, Yue, R, Zhang, R, et al. (2019) A genome-wide algal mutant library and functional screen identifies genes required for eukaryotic photosynthesis. Nature Genetics, 51: 627–635

Lin, CS, Chou, TL and Wu, JT (2013) Biodiversity of soil algae in the farmlands of mid-Taiwan. Botanical Studies, 54: 1–12

Liu, P and Lechtreck, KF (2018) The Bardet–Biedl syndrome protein complex is an adapter expanding the cargo range of intraflagellar transport trains for ciliary export. Proceedings of the National Academy of Sciences of the United States of America, 115: E934–E943

Love, MI, Huber, W and Anders, S (2014) Moderated estimation of fold change and dispersion for RNA-seq data with DESeq2. Genome Biology, 15: 1–21

Luo, M, Cao, M, Kan, Y, Li, G, Snell, W and Pan, J (2011) The phosphorylation state of an aurora-like kinase marks the length of growing flagella in Chlamydomonas. Current Biology, 21: 586–591

Luxmi, R, Blaby-Haas, C, Kumar, D, Rauniyar, N, King, SM, Mains, RE and Eipper, BA (2018) Proteases shape the Chlamydomonas secretome: comparison to classical neuropeptide processing machinery. Proteomes, 6: 36

Ma, Y, He, J, Li, S, Yao, D, Huang, C, Wu, J and Lei, M (2023) Structural insight into the intraflagellar transport complex IFT-A and its assembly in the anterograde IFT train. Nature Communications, 14: 1506

Maher Sister ML, (1947) The role of certain environmental factors in growth and reproduction of *Protosiphon botryoides Klebs*. III Reproduction. Bulletin of the Torrey Botanical Club, 74: 156–179

Mäkinen, J, Ellis, EE, Antão, LH, Davrinche, A, Laine, AL, Saastamoinen, M, Conenna, I, Hällfors, M, Santangeli, A, Kaarlejärvi, E, Heliölä, J et al. (2025) Thermal homogenization of boreal communities in response to climate warming. Proc Natl Acad Sci USA, 122: e2415260122.

Marshall, WF (2024) *Chlamydomonas* as a model system to study cilia and flagella using genetics, biochemistry, and microscopy. Frontiers in Cell and Developmental Biology, 12: 1412641

Marshall, WF and Rosenbaum, JL (2001) Intraflagellar transport balances continuous turnover of outer doublet microtubules: implications for flagellar length control. Journal of Cell Biology, 155: 405–414

May EA, Kalocsay M, D’Auriac IG, Schuster PS, Gygi SP, Nachury MV, Mick DU (2021) Time-resolved proteomics profiling of the ciliary Hedgehog response. J Cell Biol. 220: e202007207

McGoldrick, LL, Singh, AK, Demirkhanyan, L, Lin, TY, Casner, RG, Zakharian, E and Sobolevsky, AI (2019) Structure of the thermo-sensitive TRP channel TRP1 from the alga *Chlamydomonas reinhardtii*. Nature Communications, 10: 4180

Meier, HS, Schuman, IJ, Layden, TJ, Ritz, A, Kremer, CT and Fey, SB (2022) Temperature-mediated transgenerational plasticity influences movement behaviour in the green algae *Chlamydomonas reinhardtii*. Functional Ecology, 36: 2969–2982

Merchant, SS, Prochnik, SE, Vallon, O, Harris, EH, Karpowicz, SJ, Witman, GB, Terry, A, Salamov, A, Fritz-Laylin, LK, Maréchal-Drouard, et al. (2007) The *Chlamydomonas* genome reveals the evolution of key animal and plant functions. Science, 318: 245–250

Mick, DU, Rodrigues, RB, Leib, RD, Adams, CM, Chien, AS, Gyri, SP, Nachury, MV (2015). Proteomics of primary cilia by proximity labeling. Developmental Cell, 35: 497–512

Mitchell, BF, Pedersen, LB, Feely, M, Rosenbaum, JL and Mitchell, DR (2005) ATP production in *Chlamydomonas reinhardtii* flagella by glycolytic enzymes. Molecular Biology of the Cell, 16: 4509–4518

Mittag, M, Kiaulehn, S and Johnson, CH (2005) The circadian clock in *Chlamydomonas reinhardtii*. What is it for? What is it similar to? Plant Physiology, 137: 399–409

Moparthi, L, Sinica, V, Moparthi, VK, Kreir, M, Vignane, T, Filipovic, MR, Vlachova, V and Zygmunt, PM (2022) The human TRPA1 intrinsic cold and heat sensitivity involves separate channel structures beyond the N-ARD domain. Nature Communications, 13: 6113

Müller, N, Wenzel, S, Zou, Y, Künzel, S, Sasso, S, Weiß, D, Prager, K, Grossman, A, Kottke, T and Mittag, M (2017) A plant cryptochrome controls key features of the *Chlamydomonas* circadian clock and its life cycle. Plant Physiology, 174: 185–201

Murakami, H, Nobusawa, T, Hori, K, Shimojima, M and Ohta, H (2018) Betaine lipid is crucial for adapting to low temperature and phosphate deficiency in *Nannochloropsis*. Plant Physiology, 177: 181–193

Odum, M.T., Teufel, F., Thumuluri, V., Almagro Armenteros, J.J., Johansen, A.R., Winther, O., and Nielsen, H. (2024) DeepLoc 2.1: multi-label membrane protein type prediction using protein language models. Nucleic Acids Research, 52: W215–W220.

Paik, I and Huq, E (2019) Plant photoreceptors: Multi-functional sensory proteins and their signaling networks. Seminars Cell Developmental Biology, 92: 114–121

Pan, J and Snell, WJ (2005) Chlamydomonas shortens its flagella by activating axonemal disassembly, stimulating IFT particle trafficking, and blocking anterograde cargo loading. Developmental Cell, 9: 431–438

Patel, MB, Griffin, PJ, Olson, SF, Dai, J, Hou, Y, Malik, T, Das, P, Zhang, G, Zhao, W, Witman, GB and Lechtreck, KF, (2024) Distribution and bulk flow analyses of the intraflagellar transport (IFT) motor kinesin-2 support an “on-demand” model for *Chlamydomonas* ciliary length control. Cytoskeleton, 81: 586–604

Patro, R, Duggal, G, Love, MI, Irizarry, RA and Kingsford, C (2017) Salmon provides fast and bias-aware quantification of transcript expression. Nature Methods, 14: 417–419

Pazour, GJ, Agrin, N, Leszyk, J and Witman, GB (2005) Proteomic analysis of a eukaryotic cilium. Journal of Cell Biology, 170: 103–113

Pedersen LB, Geimer S, Rosenbaum JL (2006) Dissecting the molecular mechanisms of intraflagellar transport in Chlamydomonas. Current Biology, 16: 450–459

Reynolds, MJ, Phetruen, T, Fisher, RL, Chen, K, Pentecost, BT, Gomez, G, Ounjai, P and Sui, H (2018) The developmental process of the growing motile ciliary tip region. Scientific Reports, 8: 7977

Riekhof, WR, Sears, BB and Benning, C (2005) Annotation of genes involved in glycerolipid biosynthesis in Chlamydomonas reinhardtii: discovery of the betaine lipid synthase BTA1Cr. Eukaryotic Cell, 4: 242–252

Rredhi A, Petersen J, Wagner V, Vuong T, Li W, Li W, Schrader L, Mittag M (2024) The UV-A receptor CRY-DASH1 up- and downregulates proteins involved in different plastidial pathways. Journal of Molecular Biology, 436: 168271

Rredhi, A, Petersen, J, Schubert, M, Li, W, Oldemeyer, S, Li, W, Westermann, M, Wagner, V, Kottke, T and Mittag, M (2021) DASH cryptochrome 1, a UV-A receptor, balances the photosynthetic machinery of *Chlamydomonas reinhardtii*. New Phytologist, 232: 610–624

Saggiomo, M, Escalera, L, Saggiomo, V, Bolinesi, F, and Mangoni, O (2021) Phytoplankton blooms below the Antarctic landfast ice during the melt season between late spring and early summer. Journal of Phycology, 57: 541–550

Salomon, S, Schilling, M, Albrieux, C, Si Larbi, G, Jouneau, PH, Roy, S, Falconet, D, Michaud, M and Jouhet, J (2025) Betaine lipids overproduced in seed plants are not imported into plastid membranes and promote endomembrane expansion. Journal of Experimental Botany, 76: 980–996

Scaife, MA and Smith, AG (2016) Towards developing algal synthetic biology. Biochemical Society Transactions, 44: 716–722

Schmidt, M, Geßner, G, Luff, M, Heiland, I, Wagner, V, Kaminski, M, Geimer, S, Eitzinger, N, Reißenweber, T, Voytsekh, O and Fiedler, M (2006) Proteomic analysis of the eyespot of *Chlamydomonas reinhardtii* provides novel insights into its components and tactic movements. The Plant Cell, 18: 1908–1930

Schneider, CA, Rasband, WS and Eliceiri, KW (2012) NIH Image to ImageJ: 25 years of image analysis. Nature Methods, 9: 671–675

Schroda, M, Hemme, D and Mühlhaus, T (2015) The *Chlamydomonas* heat stress response. The Plant Journal, 82: 466–480

Schulze, T, Schreiber, S, Iliev, D, Boesger, J, Trippens, J, Kreimer, G and Mittag, M (2013) The heme-binding protein SOUL3 of *Chlamydomonas reinhardtii* influences size and position of the eyespot. Molecular Plant, 6: 931–944

Sharma, A, Samtani, H and Laxmi, A (2024) Molecular dialogue between light and temperature signaling in plants: From perception to thermotolerance. Journal of Experimental Botany, 75: erae356

Sharma, S, Sushmita, K, Singh, R, Sanyal, SK and Kateriya, S (2025) Phototropin localization and interactions regulates photophysiological processes in *Chlamydomonas reinhardtii*. Biochimie 239: 150–162

Snell, WJ, Eskue, WA and Buchanan, MJ (1989) Regulated secretion of a serine protease that activates an extracellular matrix-degrading metalloprotease during fertilization in Chlamydomonas. Journal of Cell Biology, 109: 1689–1694

Soneson, C, Love, MI and Robinson, MD (2015) Differential analyses for RNA-seq: transcript-level estimates improve gene-level inferences. F1000Research, 4: 1521

Trainor (1960) Mating in *Chlamydomonas chlamydogama* at various temperatures under continuous illumination. American Journal of Botany, 47: 482–484

Tuxhorn, J, Daise, T and Dentler, WL (1998) Regulation of flagellar length in Chlamydomonas. Cell Motility and the Cytoskeleton, 40: 133–146

Valledor, L, Furuhashi, T, Hanak, AM and Weckwerth, W (2013) Systemic cold stress adaptation of *Chlamydomonas reinhardtii*. Molecular & Cellular Proteomics, 12: 2032–2047

Verret, F., Wheeler, G., Taylor, A.R., Farnham, G., and Brownlee, C. (2010) Calcium channels in photosynthetic eukaryotes: implications for evolution of calcium-based signalling. New Phytologist, 187: 23–43.

von der Heyde, EL and Hallmann, A (2022) Molecular and cellular dynamics of early embryonic cell divisions in *Volvox carteri*. The Plant Cell, 34:1326–1353

Vuong, T, Shetty, P, Kurtoglu, E, Schultz, C, Schrader, L, Then, P, Petersen, J, Westermann, M, Rredhi, A., Chowdhury, S., Mukherji, R., et al. (2025) Metamorphosis of a unicellular green alga in the presence of acetate and a spatially structured three-dimensional environment. New Phytologist, 245: 1180–1196

Wemmer, K, Ludington, W and Marshall, WF (2020) Testing the role of intraflagellar transport in flagellar length control using length-altering mutants of *Chlamydomonas*. Philosophical Transactions of the Royal Society B, 375: 20190159

Wingfield JL and Lechtreck K-F (2018) *Chlamydomonas* basal bodies as flagella organizing centers, Cells 7: 79

Xiao, Y, Xin, Y, Shen, K, Yang, M, and Wu, H (2025) The roles of transient receptor potential vanilloid in autoimmune diseases. Autoimmunity Reviews, 24: 103872

Zhang, Y, Wolf-Yadlin, A, Ross, PL, Pappin, DJ, Rush, J, Lauffenburger, DA and White, FM (2005) Time-resolved mass spectrometry of tyrosine phosphorylation sites in the epidermal growth factor receptor signaling network reveals dynamic modules. Molecular & Cellular Proteomics, 4: 1240–1250

Zhang, N, Mattoon, EM, McHargue, W, Venn, B, Zimmer, D, Pecani, K, Jeong, J, Anderson, CM, Chen, C, Berry, JC, Xia, M (2022) Systems-wide analysis revealed shared and unique responses to moderate and acute high temperatures in the green alga *Chlamydomonas reinhardtii*. Communications Biology, 5: 460

Zhang, N, Venn, B, Bailey, CE, Xia, M, Mattoon, EM, Mühlhaus, T and Zhang, R (2024) Moderate high temperature is beneficial or detrimental depending on carbon availability in the green alga *Chlamydomonas reinhardtii*. Journal of Experimental Botany, 75: 979–1003

Zhao, B, Schneid, C, Iliev, D, Schmidt, EM, Wagner, V, Wollnik, F and Mittag, M (2004) The circadian RNA-binding protein CHLAMY 1 represents a novel type heteromer of RNA recognition motif and lysine homology domain-containing subunits. Eukaryotic Cell, 3: 815–825

Zou, Y, Wenzel, S, Müller, N, Prager, K, Jung, EM, Kothe, E, Kottke, T and Mittag, M (2017) An animal-like cryptochrome controls the *Chlamydomonas* sexual cycle. Plant Physiology, 174: 1334–1347

