## Supplementary Figures 1-14 for "Multi-omics studies reveal how ambient temperature changes govern cellular responses of Chlamydomonas"

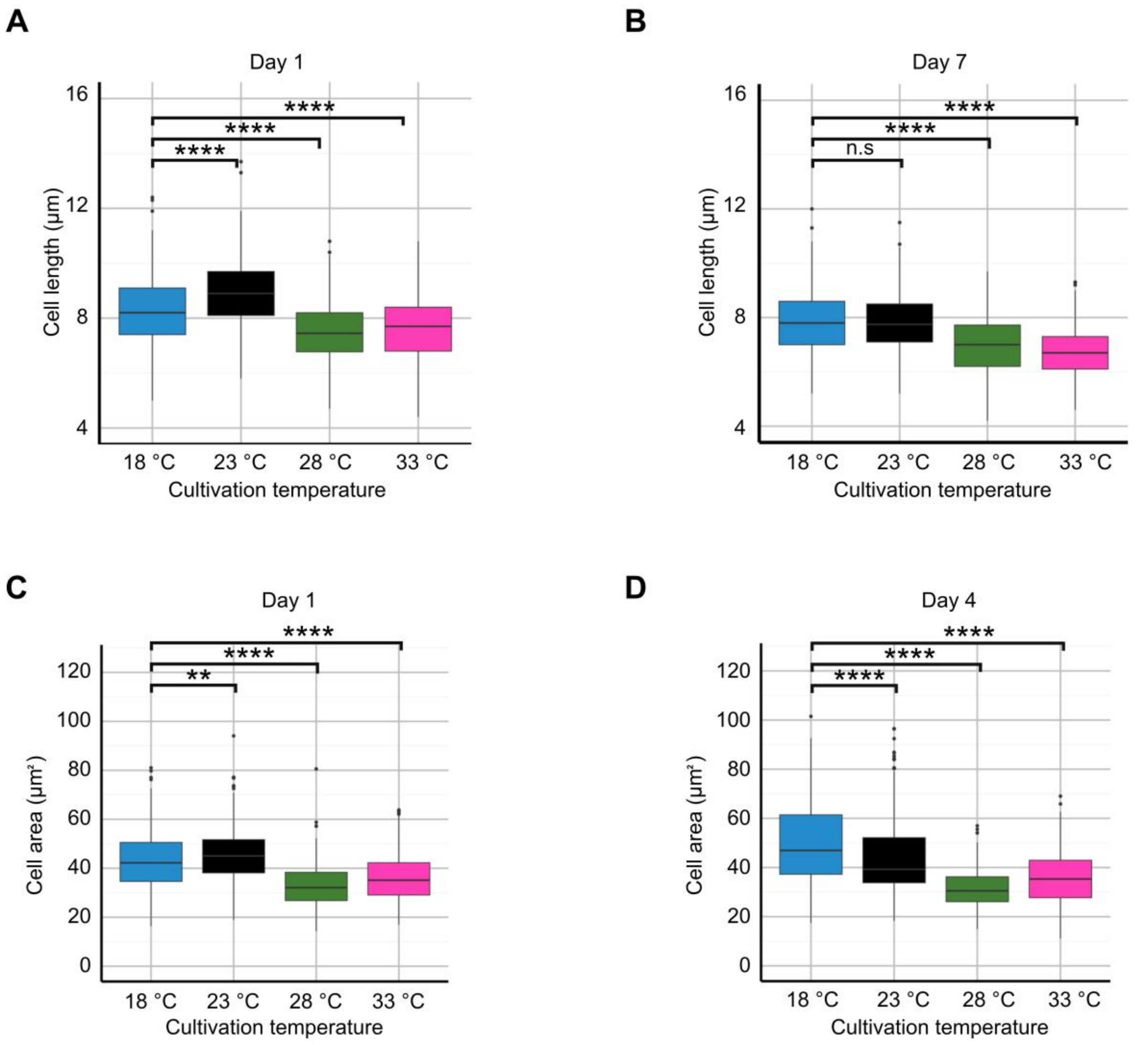


**Supplementary Figure 1. Boxplot analysis of cell lengths and estimated cell areas measured at LD6. A)** Boxplot of vertical cell length measurements (µm) on day 1 of cultivation across four temperatures (18 °C, 23 °C, 28 °C, and 33 °C). **B)** Boxplot of vertical cell length measurements (µm) on day 7 of cultivation, for the same temperature treatments. **C, D)** Boxplot of estimated cell area measurements ($\mu m^2$; see Methods) on days 1 (C) and 4 (D) of cultivation, for the same temperature treatments. **A-D)** Each boxplot represents measurements from three biological replicates (100 cells per replicate). Statistical significance between groups was determined by a two-sided *t*-test assuming unequal variance (Welch's *t*-test) and is indicated by asterisks (** adjusted *P* value <0.01; **** adjusted *P* value <0.0001; n.s.: not significant). (Support of Figure 1).

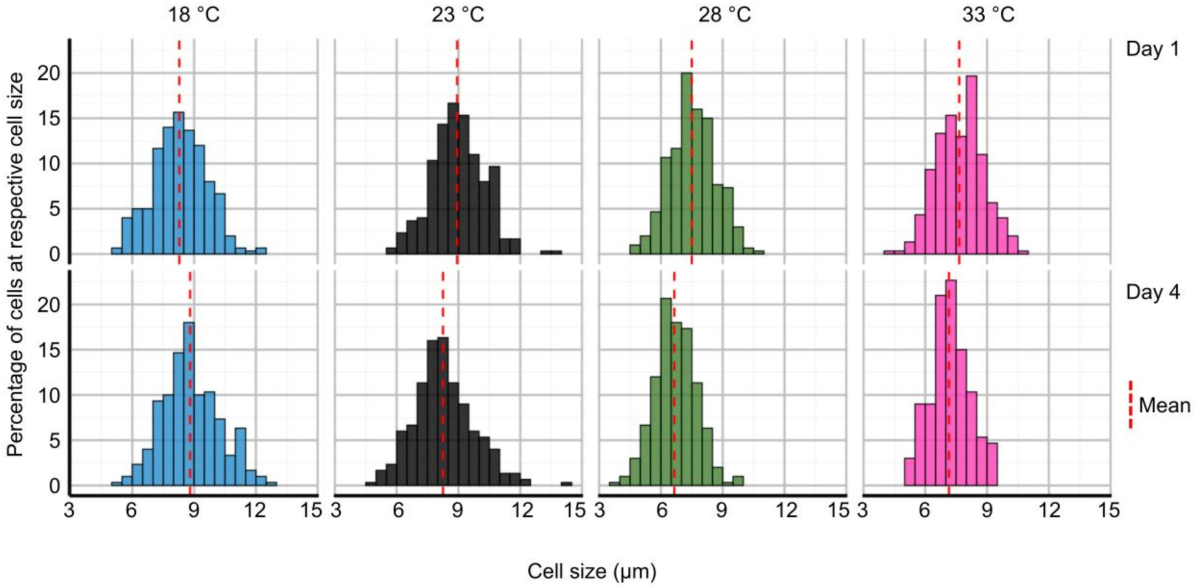


**Supplementary Figure 2. Cell length distributions at different ambient temperatures and cultivation days measured at LD6.** Frequency distribution histograms of vertical cell sizes for Chlamydomonas cultures grown at 18 °C (blue), 23 °C (black), 28 °C (green), and 33 °C (magenta). The top row of panels corresponds to measurements taken on day 1 of cultivation, and the bottom row corresponds to day 4. The dashed red line in each panel indicates the mean cell size for that specific population. This figure shows the underlying data for the boxplots presented in Figure 1B and Supplementary Figure 1A. The y-axis shows the percentage of the total cell population that falls into each specific size bin (the bars on the x-axis). For each panel, all bars add up to 100% of the measured cells for that sample. (Support of Figure 1).

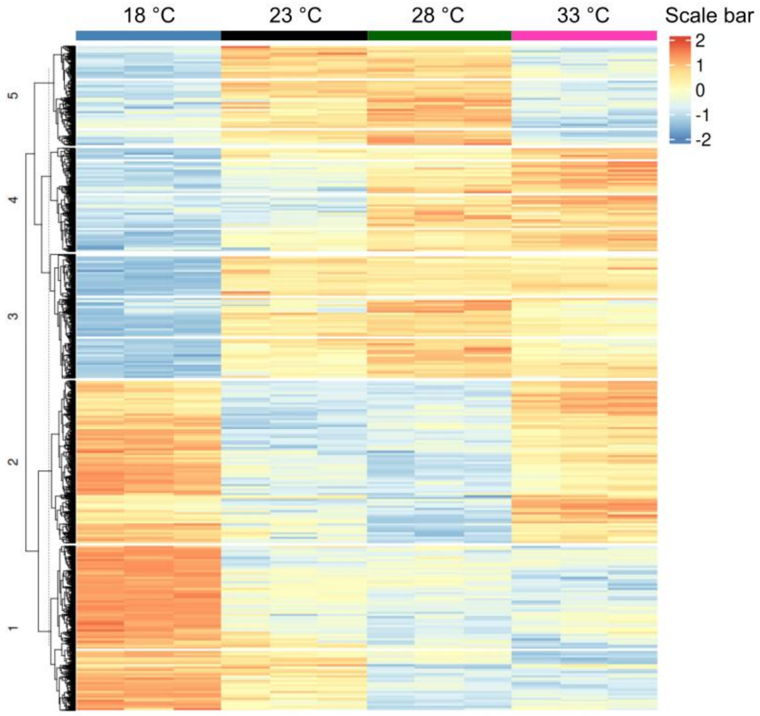


**Supplementary Figure 3. Hierarchical clustering of all differentially abundant transcripts from cells grown at 18 °C versus 28 °C.** The complete set of genes was determined using a statistical threshold of $|\log_2 FC| \geq 1$ and an adjusted *P*-value < 0.01. (Support of Figure 1).

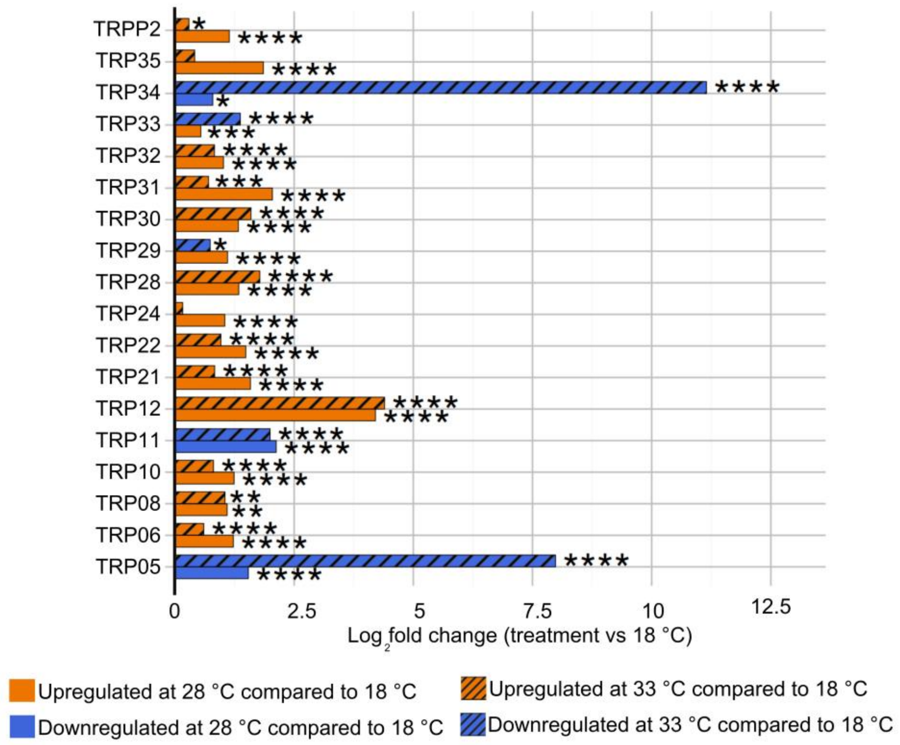


**Supplementary Figure 4. Temperature-dependent regulation of transcripts encoding TRP channels.** Barplot showing $\log_2$ fold change in the expression of all significantly changed TRP channel transcripts in Chlamydomonas in response to cultivation at varying temperatures (18 °C, 28 °C, and 33 °C). Adjusted *P*-values < 0.01 (*), < 0.001 (**), < 0.0001 (***), and < 0.00001 (****), as calculated by DESeq2, are shown. (Support of Figure 2).

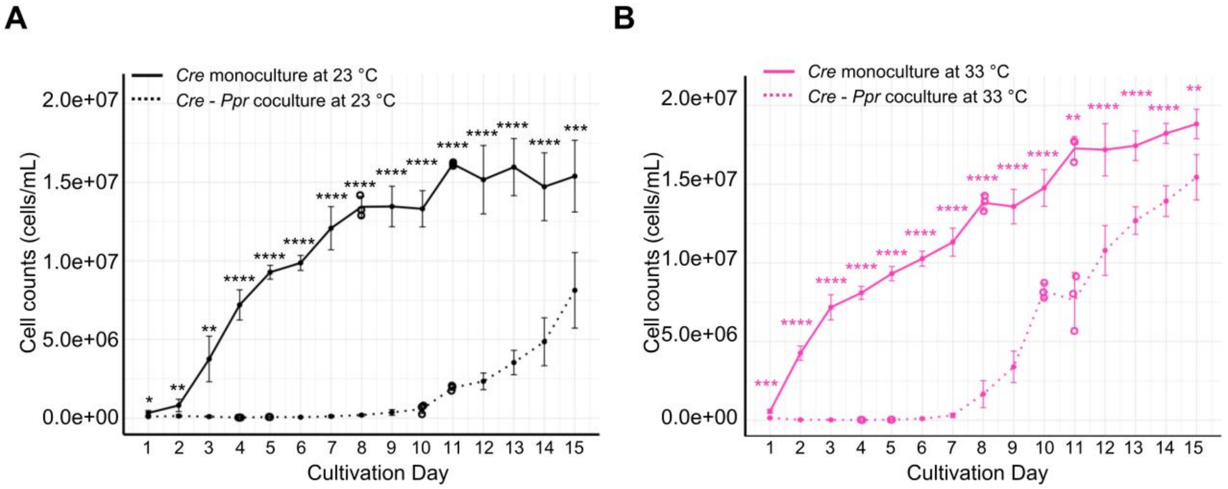


**Supplementary Figure 5. Growth curves of Chlamydomonas mono- and cocultures with *P. protegens* at 23 °C (A) and 33 °C (B).** Growth curves of axenic Chlamydomonas, abbreviated as *Cre* (monocultures), and Chlamydomonas-*P. protegens* cocultures (abbreviated as *Cre* - *Ppr*) at 23 °C and 33 °C, respectively. Cell densities (cells/mL) were measured over 15 days. Data points represent mean ± standard deviation cell densities from two independent datasets, each comprising three biological replicates, unless otherwise indicated. Data from *Cre* monocultures have been taken from Figure 1A (Supplementary Table 1). In the *Cre* - *Ppr* cocultures, only three biological replicates are available on certain days. Specifically, biological replicates are missing for days 5 and 11 in dataset 1 and for days 4 and 10 in dataset 2. Individual data from the three present biological replicates from days 4, 5, 10 and 11 are indicated by open circles. Asterisks represent significant differences estimated by a two-sided *t*-test assuming unequal variance (Welch's *t*-test). Adjusted $P$ value < 0.05 (*), adjusted $P$ value < 0.01 (**), adjusted $P$ value < 0.001 (***), and adjusted $P$ value < 0.0001 (****). (Support of Figure 2).

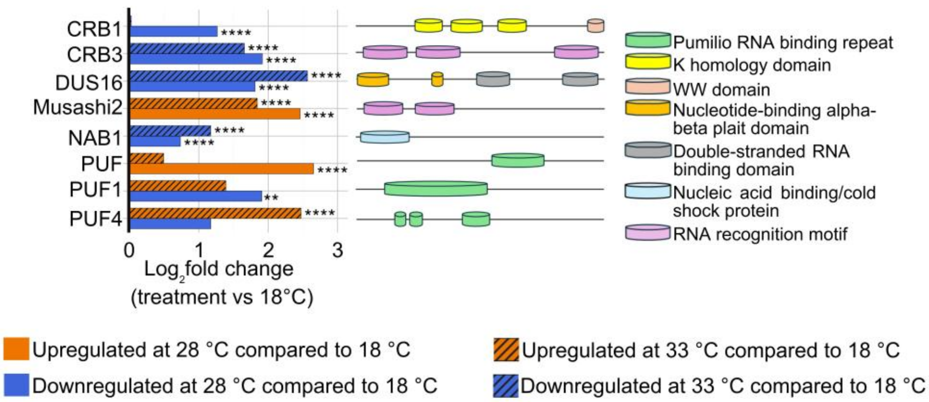


**Supplementary Figure 6. Expression profiles of transcripts encoding RNA-binding proteins.** RNA-binding proteins include Pumilio/PUF domain proteins, Musashi, CRB1/3 proteins being the C1 and C3 subunit of CHLAMY1, NAB1 (nucleic acid binding factor) and DUS16 (dull slicer-16). Known domains of the RBPs analyzed by Phytozome 14 (https://phytozome-next.jgi.doe.gov/) are highlighted. For simplicity, the size of the proteins has been standardized to a common bar length; the relative position of the domains is given. Genes are listed in Supplementary Table 5 along with their Cre accession numbers. Adjusted *P*-values < 0.001 (**) or < 0.00001 (****), as calculated by DESeq2, are shown. (Support of Figure 2).

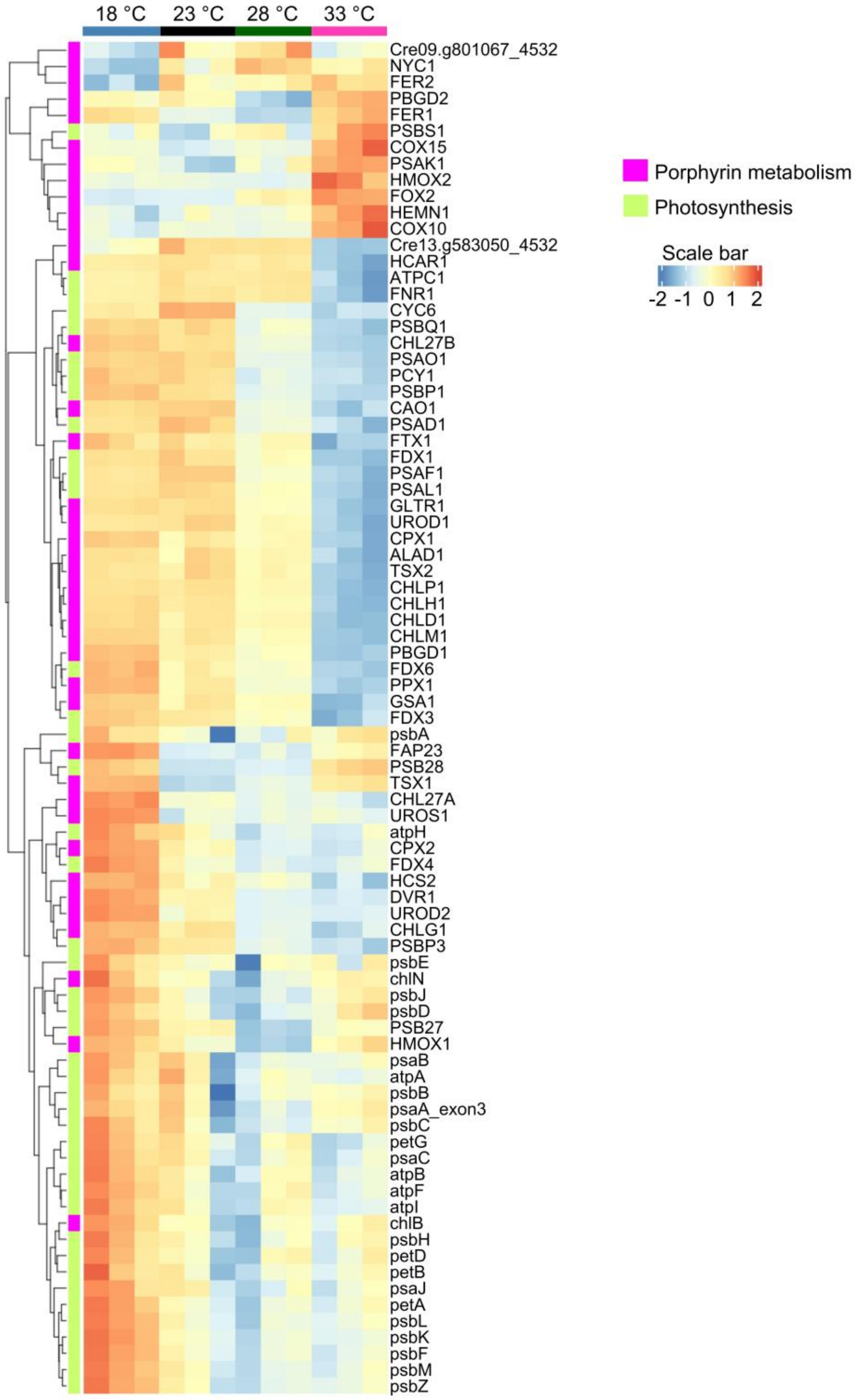


**Supplementary Figure 7. Heatmap of transcript abundances at different ambient temperatures encoding proteins of photosynthesis and porphyrin metabolism (chlorophyll biosynthesis).** Heatmap depicting expression of transcripts with a $|\log_2 FC| \geq 1$ encoding proteins of photosynthesis (KEGG pathway KO00195) and porphyrin metabolism (chlorophyll biosynthesis, KEGG pathway KO00860). Genes are listed in Supplementary Table 5 along with their Cre accession numbers. (Support of Figure 3).

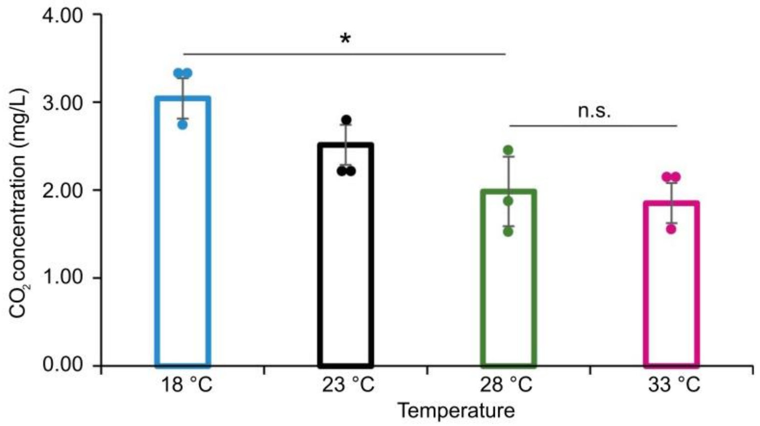


**Supplementary Figure 8: Effects of temperature on $CO_2$ levels A)** Levels of $CO_2$ in water have been measured using a carbon dioxide kit. Three biological replicates were performed (see Methods). Asterisks represent significant differences as calculated by the two-sided *t*-test assuming unequal variance (Welch's *t*-test). n.s.: not significant, *: $P < 0.05$. (Support of Figure 3).

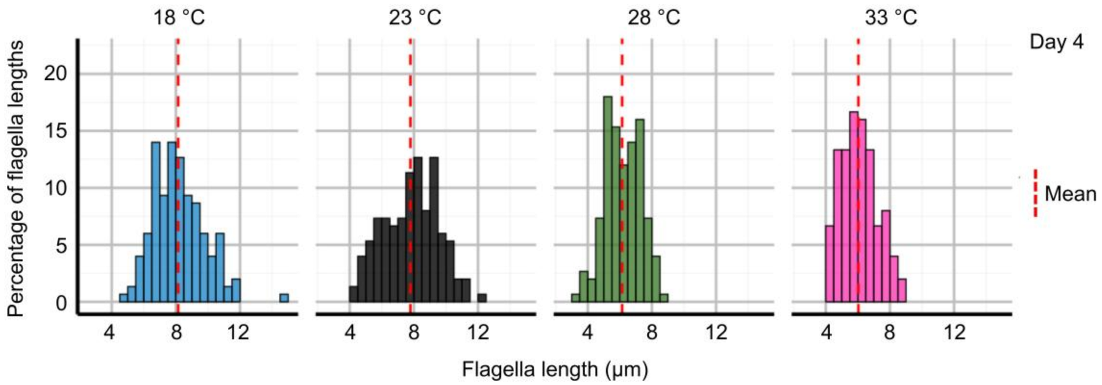


**Supplementary Figure 9. Flagellar length distributions at different ambient temperatures on day 4 at LD6.** Frequency distribution histograms of flagellar length (μm) for Chlamydomonas cultures grown at 18 °C (blue), 23 °C (black), 28 °C (green), and 33 °C (magenta). All measurements were taken on day 4 of cultivation. The dashed red line in each panel indicates the mean flagellar length for that specific population. The y-axis shows the percentage of the total flagella population that falls into each specific size bin (the bars on the x-axis). For each panel, all bars add up to 100% of the measured flagella for that sample. This figure provides the underlying data distribution for the boxplot in Figure 4D. (Support of Figure 4).

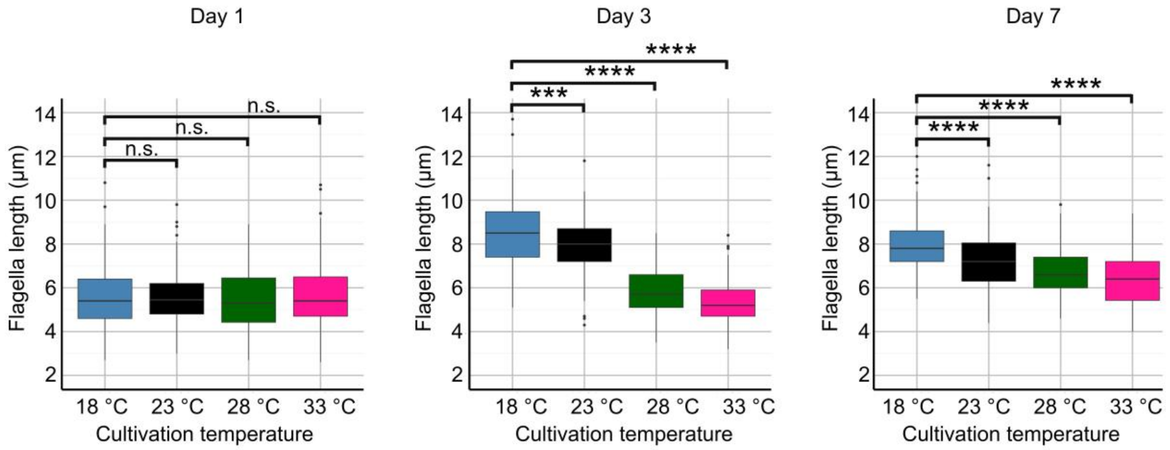


**Supplementary Figure 10. Quantification of flagellar length in cells grown at different temperatures on days 1, 3, and 7.** Quantification of flagellar lengths measured using immunolocalization images. Each boxplot includes measurements from three biological replicates (50 flagella per replicate, 150 per temperature). Significant differences between temperature treatments were determined by the two-sided *t*-test assuming unequal variance (Welch's *t*-test) and are marked by asterisks; adjusted *P* value < 0.001 (***), adjusted *P* value < 0.0001 (****), n.s. not significant. (Support of Figure 4).

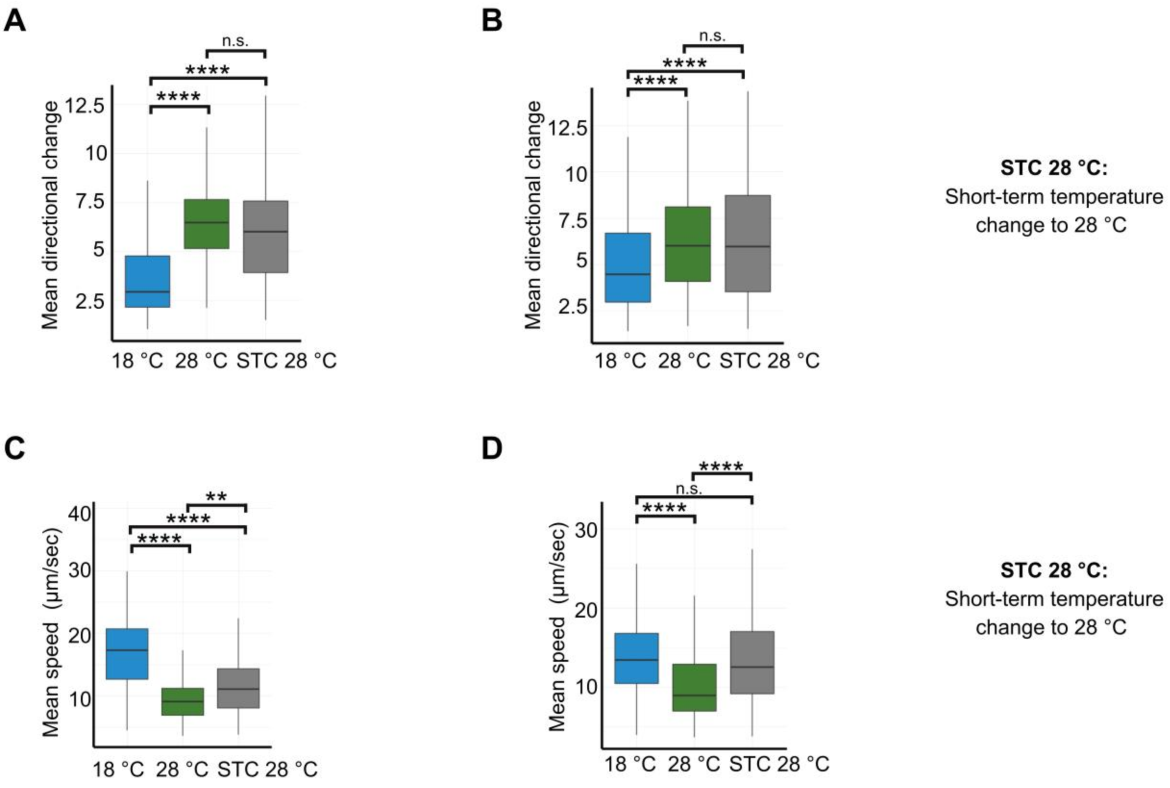


**Supplementary Figure 11. Cell movements with mean directional changes and median swimming speeds (independent replicates). A, B)** Quantification of mean directional change (degrees) for cell movements in cells grown at either 18 °C or 28 °C for four days. In one case (grey bars), cells grown at 18 °C for four days were exposed to a short-term temperature change (STC). **C, D)** Mean swimming speed (µm/sec) for cells at each temperature, calculated from tracked cell movements. **A-D)** Representative data from two different independent biological replicates for mean directional changes (A, B) and mean speed (C, D). Data reflects a minimum of 150 individual tracks per temperature per biological replicate. In B and D, a third biological replicate is present in the case of 18 °C samples. Statistical significance determined by a two-sided t-test assuming unequal variance (Welch's t-test) is indicated by asterisks (**: $P < 0.01$, ****: $P < 0.0001$, n.s.: not significant). (Support of Figure 5).

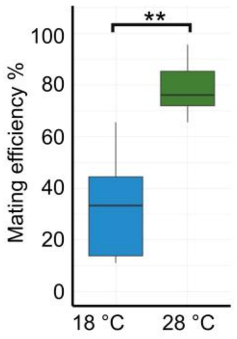


**Supplementary Figure 12. Mating efficiency at different temperatures (second independent dataset).** Quantification of mating efficiency across temperature or experimental conditions. Data represents three biological replicates with two technical replicates per biological replicate. The box plots show the proportion of successfully fused mating pairs, with mean values marked. Significant differences in mating efficiency between groups were determined using a two-sided *t*-test assuming unequal variance (Welch's *t*-test); *P*-values are indicated (**: $P < 0.01$). (Support of Figure 6).

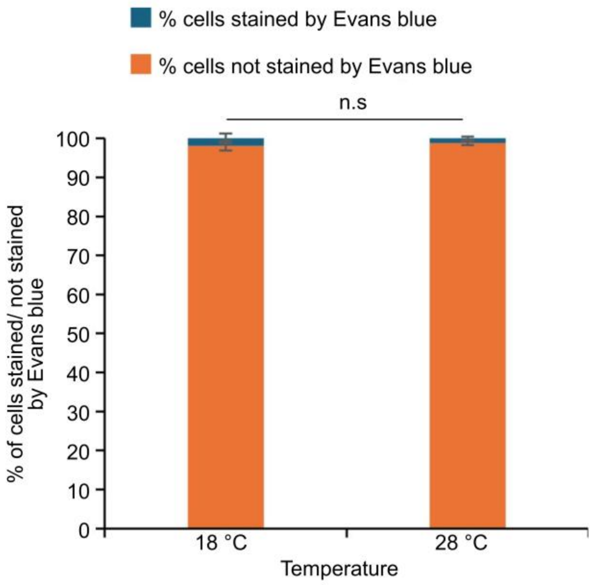


**Supplementary Figure 13. Evans Blue staining of cells grown at 18 °C or 28 °C for four days.** Evans blue staining (see Methods) was performed to determine the percentage of lysed cells that can be stained with Evans Blue (blue color) versus intact cells that cannot be stained by Evans Blue (orange color). Three biological replicates along with three technical replicates for each biological replicate were performed and the experiment was repeated twice independently. Significance was calculated by a two-sided *t*-test assuming unequal variance (Welch’s *t*-test). n.s. not significant. (Support of Figure 6).

### Codon adapted *CRY-DASH2* coding sequence within plasmid 13AA7FKP-CDS294-pMK-RQ

```
                                                 NcoI
    CACTATAGGGCGAATTGAAGGAAGGCCGTCAAGGCCGCATCCATGGAGAAAAGCCTGACC
  1 ---------+---------+---------+---------+---------+---------+
    GTGATATCCCGCTTAACTTCCTTCCGGCAGTTCCGGCGTAGGTACCTCTTTTCGGACTGG
                                                 M__E__K__S__L__T__
```

```
                         AgeI                                 BstEII
    GCAACCCATCCGCTGCGTCGTACCGGTCTGCGTACCGCACCTCTGCGTGCAACCTGGTCA
 61 ---------+---------+---------+---------+---------+---------+
    CGTTGGGTAGGCGACGCAGCATGGCCAGACGCATGGCGTGGAGACGCACGTTGGACCAGT
    A__T__H__P__L__R__R__T__G__L__R__T__A__P__L__R__A__T__W__S__
```

```
                                                  PvuII
    CCGTTTGGTGCACTGCATACCAAATTTATCGTGATTAAAAGCAGCTGCATTAGCAATAGC
121 ---------+---------+---------+---------+---------+---------+
    GGCAAACCACGTGACGTATGGTTTAAATAGCACTAATTTTCGTCGACGTAATCGTTATCG
    P__F__G__A__L__H__T__K__F__I__V__I__K__S__S__C__I__S__N__S__
```

```
    GGTGCACGTCCGGTTGCACGTCGTGCCAGCATGGGTCGTCTGCGTGCCCATGCATGTGAA
181 ---------+---------+---------+---------+---------+---------+
    CCACGTGCAGGCCAACGTGCAGCACGGTCGTACCCAGCAGACGCACGGGTACGTACACTT
    G__A__R__P__V__A__R__R__A__S__M__G__R__L__R__A__H__A__C__E__
```

```
                      BspMI     BspMI
    GCCGGTGCAGTTACCGCAGCAGGTAATGCAGGTCTGAGCAGCAGTCCGGGTCCGAGTAGT
241 ---------+---------+---------+---------+---------+---------+
    CGGCCACGTCAATGGCGTCGTCCATTACGTCCAGACTCGTCGTCAGGCCCAGGCTCATCA
    A__G__A__V__T__A__A__G__N__A__G__L__S__S__S__P__G__P__S__S__
```

```
                                                         BamHI
    CCGGTTCGTAGCCTGGGTCTGTGGTGGGTTCGTCGTGATATGCGTCTGGATGATAATGAA
301 ---------+---------+---------+---------+---------+---------+
    GGCCAAGCATCGGACCCAGACACCACCCAAGCAGCACTATACGCAGACCTACTATTACTT
    P__V__R__S__L__G__L__W__W__V__R__R__D__M__R__L__D__D__N__E__
```

```
                                                         BamHI
    GCACTGACCAGCGCAGTTCGTCATGCAGATGCAACCCTGGCAGTTCATATTCTGGATCCG
361 ---------+---------+---------+---------+---------+---------+
    CGTGACTGGTCGCGTCAAGCAGTACGTCTACGTTGGGACCGTCAAGTATAAGACCTAGGC
    A__L__T__S__A__V__R__H__A__D__A__T__L__A__V__H__I__L__D__P__
```

```
    CGTGATCTGCTGCCTCGTCGTCCGCGTAGCGAAGGTGGTCTGGGTGTTCCGAAACTGGGT
421 ---------+---------+---------+---------+---------+---------+
    GCACTAGACGACGGAGCAGCAGGCGCATCGCTTCCACCAGACCCACAAGGCTTTGACCCA
    R__D__L__L__P__R__R__P__R__S__E__G__G__L__G__V__P__K__L__G__
```

```
                                                    PvuII
    CCGCCTCGTGCGAAATTTATGCTGGAAGGTCTGAATGAACTGCGTCGTCAGCTGCAAGAA
481 ---------+---------+---------+---------+---------+---------+
    GGCGGAGCACGCTTTAAATACGACCTTCCAGACTTACTTGACGCAGCAGTCGACGTTCTT
    P__P__R__A__K__F__M__L__E__G__L__N__E__L__R__R__Q__L__Q__E__
```

```
                        PflMI           BspMI            HincII
    CTGGGTCGTGCAAGCGGTGCCCAGGCACTGGCACCAGCAGGTCCGCCTGGTCCGTCAACA
541 ---------+---------+---------+---------+---------+---------+
    GACCCAGCACGTTCGCCACGGGTCCGTGACCGTGGTCGTCCAGGCGGACCAGGCAGTTGT
    L__G__R__A__S__G__A__Q__A__L__A__P__A__G__P__P__G__P__S__T__
```

```
    GGTCCTGCGGGTGCAGCACGTGCAGAAGCAGAACTGAGCAGCGAATGTGGTGGCCTGGTT
601 ---------+---------+---------+---------+---------+---------+
    CCAGGACGCCCACGTCGTGCACGTCTTCGTCTTGACTCGTCGCTTACACCACCGGACCAA
    G__P__A__G__A__A__R__A__E__A__E__L__S__S__E__C__G__G__L__V__
```

```
    GTTCGTTATGGTCGTACCGAACAGGTTCTGCCACGTCTGCTGGGTCAGGTTCTGGAAGCA
661 ---------+---------+---------+---------+---------+---------+
    CAAGCAATACCAGCATGGCTTGTCCAAGACGGTGCAGACGACCCAGTCCAAGACCTTCGT
    V__R__Y__G__R__T__E__Q__V__L__P__R__L__L__G__Q__V__L__E__A__
```

```
                                    NdeI
     CATCCGGAACTGCGCCATGTTAGCCTGCATTATCATATGGAACCGCTGCTGGAACCGGAA
 721 ---------+---------+---------+---------+---------+---------+
     GTAGGCCTTGACGCGGTACAATCGGACGTAATAGTATACCTTGGCGACGACCTTGGCCTT
     H__P__E__L__R__H__V__S__L__H__Y__H__M__E__P__L__L__E__P__E__

     CAGGGTACAGGTGCAGGCGCAGCCGAAGATCGTGAAGCAGCAGTTCAGCGTACCGTTACC
 781 ---------+---------+---------+---------+---------+---------+
     GTCCCATGTCCACGTCCGCGTCGGCTTCTAGCACTTCGTCGTCAAGTCGCATGGCAATGG
     Q__G__T__G__A__G__A__A__E__D__R__E__A__A__V__Q__R__T__V__T__

     GCATGGGCAACCAAACAGGGTGTTGGTTGTAGCGTTCATCCGCATTGGGATAAAACCCTG
 841 ---------+---------+---------+---------+---------+---------+
     CGTACCCGTTGGTTTGTCCCACAACCAACATCGCAAGTAGGCGTAACCCTATTTTGGGAC
     A__W__A__T__K__Q__G__V__G__C__S__V__H__P__H__W__D__K__T__L__

                                             AgeI
     TATCATCCTGATGATCTGCCGTATGGTCTGTATGGCACCGGTAAAGCAGCCGGTGGTAGC
 901 ---------+---------+---------+---------+---------+---------+
     ATAGTAGGACTACTAGACGGCATACCAGACATACCGTGGCCATTTCGTCGGCCACCATCG
     Y__H__P__D__D__L__P__Y__G__L__Y__G__T__G__K__A__A__G__G__S__

                           AgeI
     GGTGGGGGTGGTGGTGGTAAAACCGGTCAGCAGCGTGCAAAACAGCAGCAAGAACGTCAA
 961 ---------+---------+---------+---------+---------+---------+
     CCACCCCCACCACCACCATTTTGGCCAGTCGTCGCACGTTTTGTCGTCGTTCTTGCAGTT
     G__G__G__G__G__G__K__T__G__Q__Q__R__A__K__Q__Q__Q__E__R__Q__

     GAACGCCACTTTACCCAGCCTGCAAATCGTGATGCACAGCGTTATCGTACCCTGCCTCCG
1021 ---------+---------+---------+---------+---------+---------+
     CTTGCGGTGAAATGGGTCGGACGTTTAGCACTACGTGTCGCAATAGCATGGGACGGAGGC
     E__R__H__F__T__Q__P__A__N__R__D__A__Q__R__Y__R__T__L__P__P__

     GTTATGACCGATTTTCGTCGTACCACCCAGAGCGCATGTGATGTTCGTGCCTGTCTGCCT
1081 ---------+---------+---------+---------+---------+---------+
     CAATACTGGCTAAAAGCAGCATGGTGGGTCTCGCGTACACTACAAGCACGGACAGACGGA
     V__M__T__D__F__R__R__T__T__Q__S__A__C__D__V__R__A__C__L__P__
```

```
                                                       PstI
     CCGCCTGCAATGCTGCCGCTGCCACCTGGTCCTTGGCGTCAGGCTGCAGCAGCAGCCGCA
1141 ---------+---------+---------+---------+---------+---------+
     GGCGGACGTTACGACGGCGACGGTGGACCAGGAACCGCAGTCCGACGTCGTCGTCGGCGT
     P__P__A__M__L__P__L__P__P__G__P__W__R__Q__A__A__A__A__A__A__

     ACAGCGGGTGGTGCCAGCGCAGCAGAAGCCGTTGCACCGGCAATGGATAGCCTGTGGGGT
1201 ---------+---------+---------+---------+---------+---------+
     TGTCGCCCACCACGGTCGCGTCGTCTTCGGCAACGTGGCCGTTACCTATCGGACACCCCA
     T__A__G__G__A__S__A__A__E__A__V__A__P__A__M__D__S__L__W__G__

                                              BspMI
     GAAATTCCGGGTAGCGTGACCGCACTGTATGAAGCCGCAGGTCCGGAAGCAGTTGCAGCT
1261 ---------+---------+---------+---------+---------+---------+
     CTTTAAGGCCCATCGCACTGGCGTGACATACTTCGGCGTCCAGGCCTTCGTCAACGTCGA
     E__I__P__G__S__V__T__A__L__Y__E__A__A__G__P__E__A__V__A__A__

     CTGGCACGGCTGCAAGAGCTGGTTGGTCTGGATTATTCACGTCTGTATCCGGGTAATCCG
1321 ---------+---------+---------+---------+---------+---------+
     GACCGTGCCGACGTTCTCGACCAACCAGACCTAATAAGTGCAGACATAGGCCCATTAGGC
     L__A__R__L__Q__E__L__V__G__L__D__Y__S__R__L__Y__P__G__N__P__

                                              AgeI      PvuII
     GCAGCAACCGCCACCGATGGCACCGCAGCAACAGCGACCGGTGGTACAGCTGCCGCAGCC
1381 ---------+---------+---------+---------+---------+---------+
     CGTCGTTGGCGGTGGCTACCGTGGCGTCGTTGTCGCTGGCCACCATGTCGACGGCGTCGG
     A__A__T__A__T__D__G__T__A__A__T__A__T__G__G__T__A__A__A__A__

      PstI                                              BspMI
     GCTGCAGCCGCATCAGATCCGCGTAGTGCATTTCCGTTTCGTGCAGGTAGTGGTGAAGCC
1441 ---------+---------+---------+---------+---------+---------+
     CGACGTCGGCGTAGTCTAGGCGCATCACGTAAAGGCAAAGCACGTCCATCACCACTTCGG
     A__A__A__A__S__D__P__R__S__A__F__P__F__R__A__G__S__G__E__A__
```

```
     CTGCGTCGCCTGCGTTATTATGTTTGGGGTAGCGCAGATTATGATCCGGAAGCGGGTGCA
1501 ---------+---------+---------+---------+---------+---------+
     GACGCAGCGGACGCAATAATACAAACCCCATCGCGTCTAATACTAGGCCTTCGCCCACGT
     L__R__R__L__R__Y__Y__V__W__G__S__A__D__Y__D__P__E__A__G__A__

     CTGGCCCTGCCGGAAGGTCAGCAACAACTGCAACAGCCGCTGGCAAGCCTGCCGAGCCTG
1561 ---------+---------+---------+---------+---------+---------+
     GACCGGGACGGCCTTCCAGTCGTTGTTGACGTTGTCGGCGACCGTTCGGACGGCTCGGAC
     L__A__L__P__E__G__Q__Q__Q__L__Q__Q__P__L__A__S__L__P__S__L__

     CTGTATTTTACCAATACCCGTGCACAGGCAGTTGGTGTTGATAGCAGCACCAAACTGAGC
1621 ---------+---------+---------+---------+---------+---------+
     GACATAAAATGGTTATGGGCACGTGTCCGTCAACCACAACTATCGTCGTGGTTTGACTCG
     L__Y__F__T__N__T__R__A__Q__A__V__G__V__D__S__S__T__K__L__S__

     CCGTTTTGTGCCCTGGGTTGTATTACACCGCGTCGTATTTATCAAGAGGTTGAAGCAGTT
1681 ---------+---------+---------+---------+---------+---------+
     GGCAAAACACGGGACCCAACATAATGTGGCGCAGCATAAATAGTTCTCCAACTTCGTCAA
     P__F__C__A__L__G__C__I__T__P__R__R__I__Y__Q__E__V__E__A__V__

     CGTGTTGCAGCAAAAGCGGCAGCCGTTCCGTGTAGCGTTGGTGCCGAAGCAGGCGGTGGC
1741 ---------+---------+---------+---------+---------+---------+
     GCACAACGTCGTTTTCGCCGTCGGCAAGGCACATCGCAACCACGGCTTCGTCCGCCACCG
     R__V__A__A__K__A__A__A__V__P__C__S__V__G__A__E__A__G__G__G__

     GGAGGTGGGGGAGCAGCACATCTGCTGCCAGCACCGGCAGAATGTGATTGGCTGGCAATG
1801 ---------+---------+---------+---------+---------+---------+
     CCTCCACCCCCTCGTCGTGTAGACGACGGTCGTGGCCGTCTTACACTAACCGACCGTTAC
     G__G__G__G__A__A__H__L__L__P__A__P__A__E__C__D__W__L__A__M__

     CATCTGTGTATTCGTGATTTTTGGACCTATACCGTTCTGAAAGAAGGTCTGGCCACCCTG
1861 ---------+---------+---------+---------+---------+---------+
     GTAGACACATAAGCACTAAAAACCTGGATATGGCAAGACTTTCTTCCAGACCGGTGGGAC
     H__L__C__I__R__D__F__W__T__Y__T__V__L__K__E__G__L__A__T__L__
```

```
     GATGAACGTGGTATTGTTGGTCAGCCGGTTAGCTGGCGTCGTGATCCTGAAGTTCTGGCA
1921 ---------+---------+---------+---------+---------+---------+
     CTACTTGCACCATAACAACCAGTCGGCCAATCGACCGCAGCACTAGGACTTCAAGACCGT
     D__E__R__G__I__V__G__Q__P__V__S__W__R__R__D__P__E__V__L__A__


     CGTTGGTGTAGCGGTCGCACAGGTCTGCCGTTTGTTGATGCAAATATGCGTGAACTGGCA
1981 ---------+---------+---------+---------+---------+---------+
     GCAACCACATCGCCAGCGTGTCCAGACGGCAAACAACTACGTTTATACGCACTTGACCGT
     R__W__C__S__G__R__T__G__L__P__F__V__D__A__N__M__R__E__L__A__


      PflMI
        AgeI                                                     BglII
     GCCACCGGTTGGATGAGCAATCGTGGTCGTCAGAATGTTGCAAGTCTGCTGGCAAAAGAT
2041 ---------+---------+---------+---------+---------+---------+
     CGGTGGCCAACCTACTCGTTAGCACCAGCAGTCTTACAACGTTCAGACGACCGTTTTCTA
     A__T__G__W__M__S__N__R__G__R__Q__N__V__A__S__L__L__A__K__D__


     PstI
     CTGCAGCAGGATTGGCGTTGGGGTGCAGAACTGTTTGAATGTCTGCTGCTGGATAGTGAT
2101 ---------+---------+---------+---------+---------+---------+
     GACGTCGTCCTAACCGCAACCCCACGTCTTGACAAACTTACAGACGACGACCTATCACTA
     L__Q__Q__D__W__R__W__G__A__E__L__F__E__C__L__L__L__D__S__D__


     GTTGCCGTGAATTATTGCAACTGGAACTATTTTGCCGGTGTTGGTAATGATCCTCGTAAT
2161 ---------+---------+---------+---------+---------+---------+
     CAACGGCACTTAATAACGTTGACCTTGATAAAACGGCCACAACCATTACTAGGAGCATTA
     V__A__V__N__Y__C__N__W__N__Y__F__A__G__V__G__N__D__P__R__N__


     CGTCGCTTTAAAACCGTTACCCAGGGAATGCAGTATGATGAAGATGCCGTTCTGGCAGCG
2221 ---------+---------+---------+---------+---------+---------+
     GCAGCGAAATTTTGGCAATGGGTCCCTTACGTCATACTACTTCTACGGCAAGACCGTCGC
     R__R__F__K__T__V__T__Q__G__M__Q__Y__D__E__D__A__V__L__A__A__


     ACCTGGCTGCCGGAACTGGCCCATCTGCCACCGCGTCTGCGTCATGCACCGTGGTCAAAT
2281 ---------+---------+---------+---------+---------+---------+
     TGGACCGACGGCCTTGACCGGGTAGACGGTGGCGCAGACGCAGTACGTGGCACCAGTTTA
     T__W__L__P__E__L__A__H__L__P__P__R__L__R__H__A__P__W__S__N__
```

```
      BspMI    PstI                                      PstI
     GACCTGCTGCCTGCAGAACAAGAGCTGGCAGCCGCAGCGACCGCAGCATCTGCAGCCTTT
2341 ---------+---------+---------+---------+---------+---------+
     CTGGACGACGGACGTCTTGTTCTCGACCGTCGGCGTCGCTGGCGTCGTAGACGTCGGAAA
     D__L__L__P__A__E__Q__E__L__A__A__A__A__T__A__A__S__A__A__F__

     AGCGCAGGCGCAACCGCACATAATGCCGGTGCCGGTATGCCGACCGTTGGCCTGGGTGGT
2401 ---------+---------+---------+---------+---------+---------+
     TCGCGTCCGCGTTGGCGTGTATTACGGCCACGGCCATACGGCTGGCAACCGGACCCACCA
     S__A__G__A__T__A__H__N__A__G__A__G__M__P__T__V__G__L__G__G__

     GAAGTTCCTGGTCCGATGGGTGCCGCAGCAGCGGCATTTGGTGTGGTTCTGGGTCGCGAT
2461 ---------+---------+---------+---------+---------+---------+
     CTTCAAGGACCAGGCTACCCACGGCGTCGTCGCCGTAAACCACACCAAGACCCAGCGCTA
     E__V__P__G__P__M__G__A__A__A__A__A__F__G__V__V__L__G__R__D__

                      BamHI
     TATCCGCTGCCGATTGTGGATCCGGTTGAACAGACCGGCACCCTGCCTGCCGAAGAAAAA
2521 ---------+---------+---------+---------+---------+---------+
     ATAGGCGACGGCTAACACCTAGGCCAACTTGTCTGGCCGTGGGACGGACGGCTTCTTTTT
     Y__P__L__P__I__V__D__P__V__E__Q__T__G__T__L__P__A__E__E__K__

                                            HindIII
     GCACGTAAAGCCGCAAAAGCCCGTGGTAAAGTTAAACTGAAGCTTCTGGGCCTCATGGGC
2581 ---------+---------+---------+---------+---------+---------+
     CGTGCATTTCGGCGTTTTCGGGCACCATTTCAATTTGACTTCGAAGACCCGGAGTACCCG
     A__R__K__A__A__K__A__R__G__K__V__K__L__K__L__

     CTTCCTTTCACTGCCCGCTTTCCAG
2641 ---------+---------+-----
     GAAGGAAAGTGACGGGCGAAAGGTC
```

**Supplementary Figure 14: Codon adapted *CRY-DASH2*.** The *CRY-DASH2* coding sequence was codon-adapted for *E. coli* and synthesized by GeneArt within plasmid 13AA7FKP-CDS294-pMK-RQ. (Support for the Methods section “Crude extracts and immunoblots”).
